# Variable performance of widely used bisulfite sequencing methods and read mapping software for DNA methylation

**DOI:** 10.1101/2025.03.14.643302

**Authors:** Emily V. Kerns, Jesse N. Weber

## Abstract

DNA methylation (DNAm) is the most commonly studied marker in ecological epigenetics, yet the performance of library preparation strategies and bioinformatic tools are seldom assessed in genetically variable natural populations. We profiled DNAm in threespine stickleback (*Gasterosteus aculeatus*) liver tissue, using reduced representation bisulfite sequencing (RRBS) and whole genome bisulfite sequencing (WGBS) across technical and biological replicates. We additionally collated publicly available RRBS and WGBS data from taxonomically diverse organisms, and then compared how the most commonly used methylation software (Bismark) performed relative to alternative pipelines (BWA meth, BiSulfite Bolt, and Biscuit). Even after choosing parameters to maximize Bismark’s mapping efficiency, it was still outperformed by all other methods. Surprisingly, newer tools overrepresented DNAm compared to older methods, highlighting the importance of testing methods on nonmodel organisms. There were also distinct differences in DNAm profiles produced across library preparation methods, with large impacts of population and read depth filters. Methylated sites unique to WGBS predominantly mapped to introns and intergenic regions, while sites unique to RRBS primarily overlapped with promoters and exons. Moreover, the prevalence of nucleotides with intermediate methylation (within individuals) was greatly reduced in RRBS. Together, this suggests that RRBS may be more useful for detecting functionally-relevant methylation differences. Based on these results, we provide methodological recommendations for improving the reliability and utility of DNAm profiles, particularly concerning the detection of functionally relevant DNAm differences in genetically diverse natural populations.

## Introduction

Epigenetics is an umbrella term for chemical modifications to DNA and proteins, with DNA methylation (DNAm) being among the most well studied epigenetic marks in ecologically focused studies (Banta & Richards, 2018; Hu & Barrett, 2017; Lea et al., 2018; Rey et al., 2020). As molecular ecologists increasingly explore the role of methylation in natural systems, there is a need for increased attention to how methodological choices impact inferences of between and within individual variation. This is particularly important given that most methylation profiling methods were originally developed and applied in inbred laboratory model organisms.

There are multiple forms of DNAm, but the most abundant type in vertebrates is the addition of a methyl group to cytosine at CpG dinucleotides (Jaenisch & Bird, 2003). DNAm has a broad range of genomic effects, including silencing transposable elements (Marin et al., 2020), blocking transcription factors and other proteins from binding to DNA, increasing the likelihood of spontaneous deamination of methylated cytosines converting them to thymine (Ord et al., 2023), and slowing down transcriptional machinery (Cholewa-Waclaw et al., 2019). While its role varies, increased methylation is often associated with decreased gene expression (Zemach et al., 2010). Multigenerational common garden experiments show that while DNAm profiles are largely heritable they can also be modified by environmental stimuli (Bogan et al., 2023; Brennan et al., 2025; Heckwolf et al., 2020; Hu et al., 2021). Moreover, environmentally-induced shifts in DNAm can be inherited across generations (Takahashi et al., 2023; Tobi et al., 2018). Recent findings also suggest that methylated regions experience different selection pressures than background genomic DNA (Brennan et al., 2025; Ord et al., 2023). Despite accumulating evidence that DNAm may play an important role in evolution, this topic is still an active area for debate (Adrian-Kalchhauser et al., 2020; Charlesworth et al., 2017; Jaenisch & Bird, 2003; Otterdijk & Michels, 2016). The increasing interest in understanding functional and evolutionary roles of DNAm warrants greater focus on how methodological choices affect the detection and quantification of DNAm.

DNAm is commonly measured using bisulfite sequencing methods (BS-seq). Treating DNA with bisulfite converts unmethylated cytosines to uracil, which then becomes thymine during PCR amplification. When compared to a reference genome, C/T mismatches are identified as unmethylated cytosines, while any cytosines remaining are inferred to be methylated. Although DNAm at any particular base pair is a binary trait (i.e., present/absent at a CpG site on each read), the same location may not be methylated across cells or tissues. Some CpG sites will be consistently methylated (or unmethylated) while others will have intermediate methylation (Hay et al., 2023; Wang et al., 2020). When BS-seq is used for bulk sequencing of many cells, the three possible methylation states of a CpG site in a single cell (present/absent/heterozygous) are summed and become a quantitative metric (i.e., proportion of total sites that are methylated). Accurately measuring this quantitative variation is contingent on the number of times that a given base pair is represented in the sequencing data, often referred to as read depth.

The two most commonly used BS-seq methods are whole genome (WGBS) and reduced representation bisulfite sequencing (RRBS; Gu et al., 2011). While WGBS offers the potential to profile all CpG sites, it requires large amounts of sequencing data in order to repeatedly sample individual DNAm sites. This requirement often limits WGBS to experiments involving few biological replicates and relatively low read depth, which reduces accuracy of methylation calls and statistical power to detect group-level differences (Seiler Vellame et al., 2021). Additionally, because WGBS does not enrich for CpG sites, much of the sequence data is uninformative for epigenetic studies. For example, 70-80% of mapped reads in human WGBS studies do not contain CpG dinucleotides (Elliott et al., 2015). In contrast, RRBS uses methylation-insensitive restriction enzymes to target sequencing toward CpG islands (e.g., the most commonly used enzyme is MspI, with a cut site of CC/GG). CpG islands are regions of the genome with a high density of CpG sites and believed to be functional hotspots for DNAm (Suzuki & Bird, 2008). Thus, RRBS not only enriches for regions most likely to harbor functionally important methylation differences, but also by profiling <10% of the genome researchers can increase sample sizes and read depth of regions covered (Bock et al., 2010; Laine et al., 2022). The large differences in the types of data recovered from WGBS (wide breadth of the genome sequenced, lower read depth, and smaller sample sizes) and RRBS (specifically targets CpG islands, higher read depth, and larger sample sizes) make each approach better for particular types of research questions. Because evolutionary and ecological studies often assess group-level variation, and therefore require relatively large sample sizes, RRBS is becoming the preferred epigenetic profiling method for these disciplines. However, few RRBS studies have directly compared its results with WGBS, making it difficult to directly assess what biases or limitations are inherent to each method.

Apart from the differences in library construction and experimental design, DNAm also requires specialized analysis methods. For example, bioinformatic tools apply different approaches to account for unmethylated cytosines no longer matching the reference genome after bisulfite treatment. Bismark (Krueger & Andrews, 2011), the most common tool for mapping bisulfite converted sequences (Laine et al., 2022) (Table 1), handles mismatches by producing *in silico* sense (C → T) and antisense (G → A) conversions at all CpG sites before aligning reads to a reference genome that is itself converted using the same process. Reads are considered ambiguous and discarded if they align with multiple versions of the *in silico* converted reference genome. However, this software relies on Bowtie2 (Langmead & Salzberg, 2012) for read mapping, which often produces lower mapping efficiency (percentage of reads mapped to the reference) than other commonly used mapping algorithms such as BWA mem (Harrath et al., 2019). Notably, BWA mem uses local alignment, meaning that if a small portion of a sample read does not align well to the reference genome it is clipped for a small penalty. Meanwhile, Bowtie2 requires global alignment, requiring the entire read to successfully map without additional trimming. Tuning Bismark mapping parameters can improve mapping efficiency but may pose a substantial hurdle for novice users. Additionally, generating four intermediate *in silico* conversions for both strands of the reference genome and sample reads leads to longer computational run times and greater memory demands than alternative tools (Farrell et al., 2021; Krueger & Andrews, 2011; Nunn et al., 2021).

**Table 1:** Search results for different bioinformatic tools associated with bisulfite converted DNA. The search terms used were “software name” AND “DNA methylation,” conducted on September 8, 2025.

| Tool | Google Scholar Search Results | Web of Science Search Results |
| --- | --- | --- |
| **Bismark** | 7,110 | 36 |
| **BWA meth** | 396 | 8 |
| **BiSulfite Bolt** | 74 | 1 |
| **Biscuit** | 372 | 5 |
| **MethylDackel** | 479 | 1 |

While Bismark remains the most popular BS-seq aligner, there are a number of alternative software wrappers that use the BWA mem algorithm. BWA meth (Pedersen et al., 2014), the oldest such wrapper, is faster than Bismark because it only performs *in silico* conversion of the reference genome prior to read mapping, not the sampled reads (Nunn et al., 2021; Sun et al., 2018). However, a current limitation of BWA meth is that it stops after mapping reads, recommending users rely on MethylDackel (https://github.com/dpryan79/MethylDackel) to tabulate methylation. BiSulfite Bolt (Farrell et al., 2021) and Biscuit (Zhou et al., 2024) are more recent aligners that also perform methylation calling. Both BiSulfite Bolt and Biscuit create a 3-base BWA-indexed reference genome that has been bisulfite converted *in silico* (Farell et al., 2021; Zhou et al., 2024). However, Biscuit is unique in that it also creates a 4-base indexed reference and uses asymmetrical alignment to increase mapping efficiency of unmethylated cytosines. Biscuit will map T’s (bisulfite-converted unmethylated cytosines on the forward strand) and A’s (bisulfite-converted unmethylated cytosines on the reverse strand) from the sample read to C’s and G’s in the reference genome, but a G and C in the sample cannot be aligned with an A and T in the reference. These methods have yet to be thoroughly compared across a diverse set of wild organisms.

When working with BS-seq data from genetically variable organisms, C/T mismatches between the sample read and reference can result from either an unmethylated cytosine or a nucleotide polymorphism. While there are a number of available tools to filter SNPs from BS-seq data (Lindner et al., 2022), here we focus on MethylDackel (https://github.com/dpryan79/MethylDackel). MethylDackel allows the user to specify the maximum proportion of non-Gs permitted on the opposite strand before a site is considered a SNP and excluded from the dataset. Notably, depth influences the accuracy of SNP filtering with this method, which may be constrained if higher depth was sacrificed for increased genomic breadth (RRBS versus WGBS). Importantly, SNP filtering may impact estimates of DNAm variation if polymorphic sites are frequently eliminated from analyses. This type of filtering may be particularly important for quantifying DNAm in natural populations where individual genotypes are unknown.

Here, we provide a comparison of approaches for generating and analyzing DNAm data that are particularly relevant to molecular ecologists. Specifically, we generated WGBS and RRBS data from five ecologically-divergent populations of threespine stickleback (*Gasterosteus aculeatus*). Four individuals were profiled using both WGBS and RRBS to assess how the methods diverge in: 1) which and how many DNAm sites are recovered, 2) depth of coverage, and 3) distributions of percentage DNAm per site. We then compared the results of several bioinformatic pipelines when applied our stickleback dataset and five other taxonomically-diverse organisms (cichlid (*Astatotilapia calliptera*), purple sea urchin (*Strongylocentrotus purpuratus*), great tit (*Parus major*), coral (*Acropora nana*), and stick insect (*Timema cristinae*)). While studies have assessed performance of bioinformatic pipelines for human and plant methylation data (Farrell et al., 2021; Nunn et al., 2021; Sun et al., 2018; Zhou et al., 2024), to our knowledge this has not yet been done in a comparative context that considers natural animal populations despite taxonomic variation in genome-wide methylation patterns (Klughammer et al., 2023). We predicted that RRBS would provide fewer total DNAm sites, greater per-site read depth, and that most DNAm would be found in CpG islands (Bock et al., 2010). We also predicted that BWA mem-based read mappers would perform better than Bowtie-based mappers in terms of total mapping efficiency. After explaining our findings, we discuss how to optimize sequencing resources for ecological epigenetics, particularly for systems where DNA polymorphisms are unknown. Finally, we consolidate information for each of our computational wrappers for bisulfite alignment, providing open access code to encourage researchers to test various read mapping methods, optimize their own bioinformatic pipelines, and increase the availability of data for downstream analysis.

## Methods

### Sample and Data Collection

We sampled liver tissue from wild and lab-reared threespine stickleback across five populations (Supplemental Table 1; UW-Madison IACUC protocol L006460-A01; Alaska Department of Fish & Game permit number SF2022-043, BC Ministry of Forests permit number NA23-787881). RRBS was applied to all samples (n=34), while WGBS (n = 4) was used for common garden fish from two Alaskan populations: Watson and Wik Lakes. We extracted DNA from each sample using carboxyl coated magnetic beads (BOMB.bio method: Oberacker et al., 2019). Bisulfite conversion, library preparation, and sequencing were performed by Admera Health Biopharm Services. Some fish were exposed to the tapeworm *Schistocephalus solidus* as part of a separate study. None of these individuals displayed obvious differences in DNAm patterns, and therefore were retained for further analysis (Supplemental Figure 1). Notably, each barcoded RRBS sample was mixed into one of two sequencing pools (Supplemental Table 1), which produced an average (± SEM) of 10.5 (± 1.3) million and 31.5 (± 15.4) million reads per sample following adapter trimming. Given that CpG islands comprise of ∼8.5% of the threespine stickleback genome (see *Genomic Annotations of RRBS-WGBS Technical Replicates* for CpG island definition), this is predicted to result in a depth of 20.7x and 62.2x for batch 1 and 2, respectively. Based on PC1 from a PCA (methylKit v.1.33.3, Akalin et al., 2012) separated samples by sequencing batch, but batch effects dissipated along PC2 (Supplemental Figure 2). All WGBS samples were sequenced in one batch, resulting in 58.5 (± 1.0) million reads per sample after adapter trimming, which is predicted to result in an average depth of 9.8x.

In addition to generating our own data from stickleback, we used publicly available data from five other animals to assess the performance of BS-seq bioinformatic pipelines (Supplemental Table 2). Our four stickleback samples were the only data we could find in which the same samples were used for both RRBS and WGBS. We gathered RRBS data from a cichlid (*Astatotilapia calliptera*; n = 33; Vernaz et al., 2021), great tit (*Parus major*; n = 61; Viitaniemi et al., 2019), and purple sea urchin (*Strongylocentrotus purpuratus*; n = 12; Bogan et al., 2023). We used WGBS data from a cichlid (*Astatotilapia calliptera*; n = 6; Vernaz et al., 2021), coral (*Acropora nana*; n = 30; Guerrero & Bay, 2024), and stick insect (*Timema cristinae*; n = 7; de Carvalho et al., 2023). All of the publicly available RRBS datasets were single end reads, while the WGBS data were paired end reads.

### Read Mapping

We applied the same bioinformatic pipelines to all datasets. First, we used the -q 20 option in Trim Galore! v. 0.6.10 (https://github.com/FelixKrueger/TrimGalore) to trim adapters and barcodes, also using the -paired and -rrbs options (where appropriate) to remove cytosines filled in at the end of reads as an artifact of library preparation. We then aligned reads to the reference genome of each organism (Supplemental Table 2) using default parameters in four programs: BWA meth v. 0.2.7, Bismark v. 0.24.2, BiSulfite Bolt v. 1.4.8, and Biscuit v. 1.7.0. Bismark resulted in a low mapping efficiency 28.10 (± 2.26%) for the RRBS stickleback samples, so we followed the guidance in the documentation and aligned the data in single-end mode with the flag -nondirectional, resulting in a mapping efficiency analogous to other studies. As Bismark performs global alignment while all of the other tools perform local alignment, we also used the -local option to have Bismark perform local alignment (“Bismark Local”), making it more comparable to the other methods. For BWA meth, BiSulfite Bolt, and Biscuit alignments, samtools v. 1.9 (Danecek et al., 2021) was used to convert SAM outputs to BAM format. We used Bamtools v.2.5.2 (Barnett et al., 2011) to calculate mapping efficiency for each method except Bismark, which includes mapping efficiency in the output. For all other methods, we calculated mapping efficiency as the percentage of reads mapped to the genome - the percent of mapped reads that failed QC.

### Methylation and SNP Calling

We sought to understand how choice of read mapping method may influence downstream methylation profiles. We therefore used MethylDackel v. 0.5.1 (https://github.com/dpryan79/MethylDackel) to call methylation and filter SNPs from the threespine stickleback bam files produced with BWA meth, BiSulfite Bolt, and Biscuit, even though BiSulfite Bolt and Biscuit have their own methylation calling software. Methylation call accuracy scales logistically with read depth and becomes asymptotic at 10x (Seiler Vellame et al., 2021), so we only retained sites that were sequenced at least 10 times. It can be difficult to differentiate between PCR replicates and true biological duplicates in RRBS data due to biased genomic fragmentation (Laine et al., 2022). Therefore, for the RRBS samples only, we followed recommendations in the MethylDackel documentation and applied the --keepDupes option. To prioritize capturing methylation variation, we used a lenient SNP filter (--maxVariantFrac 0.8, -- minDepth 5) in MethylDackel. The final methylation file was modified to be compatible as an input to methylKit in R (Akalin et al., 2012). Bismark’s bam files were incompatible with MethylDackel, so methylation calls were performed using the standard Bismark algorithm.

### Statistical analyses of BS-seq Alignment Wrappers

We applied a number of statistical tests in R v. 4.3.2 (R Core Team, 2023) to compare read mapping approaches with both RRBS and WGBS data. We used the nonparametric Friedman test of repeated measures to assess whether the mapping efficiency of individual libraries differed between alignment software. Effect sizes were calculated using Kendall’s W, with scores of 0.1-0.3 indicating small effect, 0.3-0.5 moderate effect, and >0.5 is a large effect Post hoc pairwise comparisons were done using a Wilcox signed-rank test, with p-values corrected for multiple comparisons using the Bonferroni method in rstatix v. 0.7.2 and coin v.1.4-3 (Hothorn et al., 2008; Kassambara, 2023).

To determine if the resulting methylation profiles of individuals were in agreement despite differences in read mapping approach, we created a linear model in ggpmisc v. 0.6.1 (Aphalo, 2024) to compare the mean percent methylation obtained using each pipeline. A slope near 1 and intercept near zero indicates high agreement between pipelines. We then visualized the relationship between the residuals of each model and per-sample sequencing depth, predicting that agreement between software would increase with depth.

### Genomic Annotations of RRBS-WGBS Technical Replicates

We next assessed differences between RRBS and WGBS results in stickleback. We tabulated the proportion of CpG sites that were shared either (1) between methods within an individual (i.e. technical replicates; Supplemental Table 3) or (2) between individuals using the same method (i.e. biological replicates; Supplemental Table 4). As we were interested in the breadth of genomic coverage between library preparation methods, we did not filter SNPs prior to this comparison and relaxed the minimum depth (tested at both 5x and 10x). The frequency distribution of percent methylation per base was used to visualize how the overall methylation profile for an individual varied based on the library preparation method.

To determine where RRBS and WGBS vary in their coverage of the genome, we conducted genomic annotations on CpG sites that were sequenced with both methods, RRBS only, and WGBS only with a minimum depth of 5x. TaJoCGI (Nell, 2020) identified CpG islands in the v.5 threespine stickleback reference genome (Nath et al., 2021) based on the algorithm from Takai & Jones (2002). We quantified the overlap between methylation sites and CpG islands and shores (±2 kbp CpG islands) with genomation v. 1.34.0 (Akalin et al., 2015). We next assessed differences in functional regions captured by the two BS-seq methods. We specifically examined variation in 1) the proportion of CpG sites that overlapped with promoters, gene bodies, and intergenic regions and 2) the proportion of all promoters and gene bodies in the genome that were captured. For more information regarding library preparation, sequencing, and analysis, refer to the supplemental methods.

## Results

### Read Mapping: BWA mem-based algorithms outperform Bismark

The five sequence alignment tools produced significantly different mapping efficiency (RRBS: Friedman test p_adj_ = 1.30 x 10^−60^, df = 4, Kendall’s W = 0.514, Figure 1A; WGBS: Friedman test p_adj_ = 3.04 x 10^−39^, df = 4, Kendall’s W = 0.992, Figure 1B). The most consistent pattern was that local mapping outperformed Bismark’s default method of global alignment. Simply running Bismark with the “-local” parameter greatly increased mapping efficiency (RRBS Bismark - Bismark Local: p_adj_ = 1.38 x 10^−23^, effect size r = 0.868; WGBS Bismark - Bismark Local: p_adj_ = 2.47 x 10^−8^, effect size r = 0.871). Despite this increase in performance, BWA mem-based aligners tended to result in higher mapping efficiency than Bismark Local. For example, nearly 100% of the Cichlid sample reads were mapped to the reference using Biscuit and BWA meth (RRBS Biscuit: 99.5 ± 0.103% (SD), RRBS BWA meth: 98.7 ± 0.564%), compared to a mean mapping rate of 85.2 ± 1.56% RRBS reads with Bismark Local. Meanwhile, BiSulfite Bolt was clearly the best method for Great Tits in terms of mapping efficiency (93.0 ± 2.58%), with a mapping rate almost 12% higher than the second best aligner (Great Tit BiSulfite Bolt - Bismark Local: p_adj_ = 1.38 x 10^−23^, effect size r = 0.870). Overall, Biscuit displayed the highest mapping efficiency, closely followed by BiSulfite Bolt (RRBS Bisulfite Bolt - Biscuit: p_adj_ > 0.05, effect size r = 0.220; WGBS: p_adj_ = 1.85 x 10^−9^, effect size r = 0.804) and BWA meth (RRBS Biscuit - BWA meth: p_adj_ = 1.24 x 10^−6^, effect size r = 0.444; WGBS: p_adj_ = 1.42 x 10^−13^, effect size r = 0.871).

**Figure 1:**
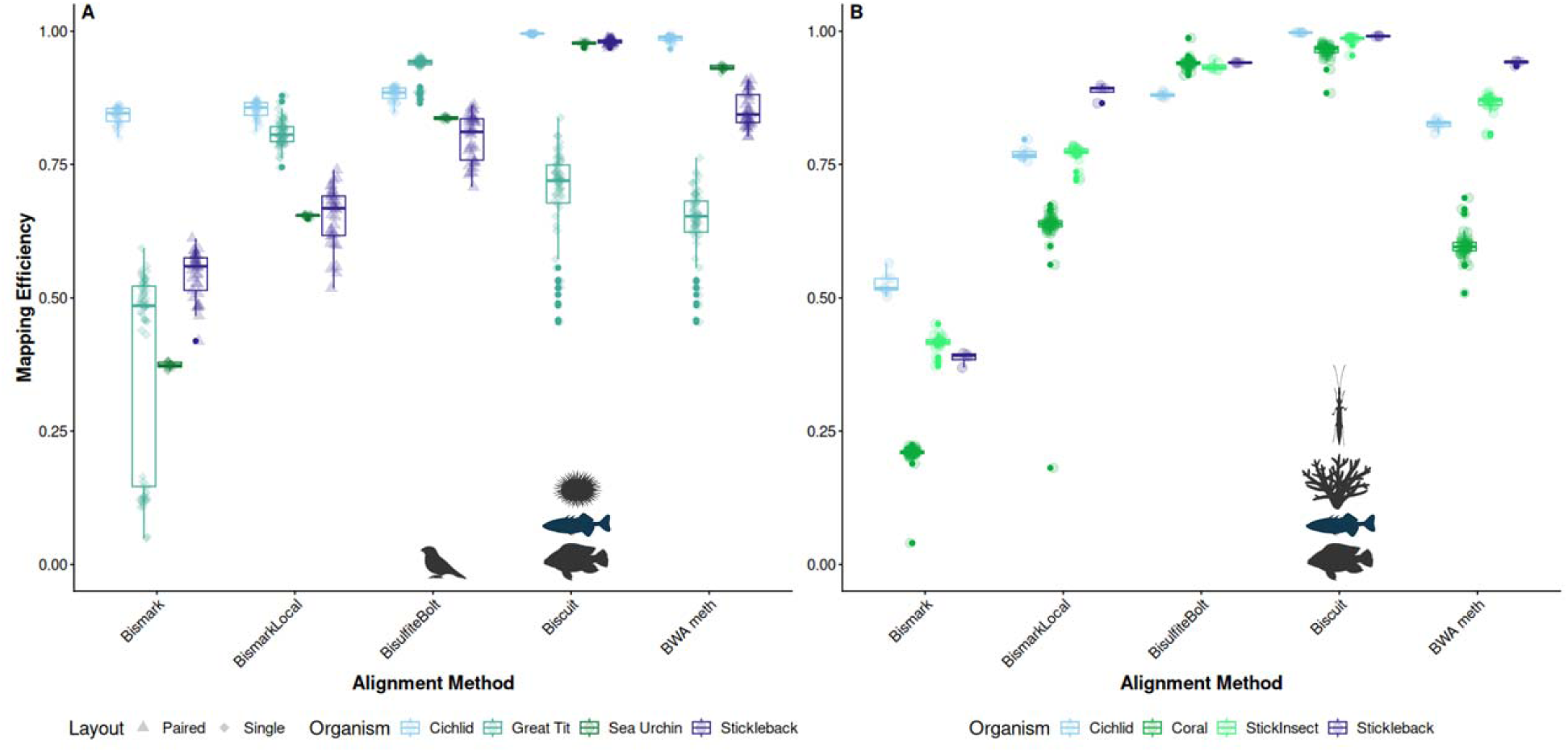
Interspecific variation in mapping efficiency of five alignment tools using **(A)** RRBS and **(B)** WGBS data. Layout refers to whether samples were sequenced using paired end or single end reads. All WGBS data was sequenced with paired end read chemistry. Symbols (Biorender) denote the read mapping software that resulted in the highest mapping efficiency for each organism. Threespine stickleback are the dark blue fish, while cichlid are the dark gray fish.

Because a PCA revealed clear sequencing batch effects among the stickleback samples (Supplemental Figure 2), we assessed whether variation in mapping efficiency across alignment tools was connected to batch effects. Although Bismark’s mapping efficiency did not differ between sequencing batches (Wilcox rank sum exact test, W = 142, p-value = 0.732), there were small but significant batch effects for every other BS-seq aligner (Supplemental Figure 3).

### Read mapping software & sequencing effort influence reliability of downstream methylation calls

We next tested if read mapping methods produced similar methylation profiles in threespine stickleback sequenced with RRBS despite significant differences in mapping efficiency. While Bismark resulted in the lowest mapping efficiency among all read aligners we tested (54.4 ± 4.35%), the average percent methylation of each individual was remarkably similar to that produced from BWA meth, which had an average mapping efficiency of 85.3 ± 3.23% (slope = 0.935, intercept = 9.24; Figure 2A). Next, because Bismark Local had a higher mapping efficiency than Bismark (65.2 ± 5.55% versus 54.4 ± 4.35%, respectively; p_adj_ = 5.63 x 10^−6^, effect size r = 0.873), we compared the average percent methylation of each sample recovered from these two methods, resulting in highly similar methylation profiles (slope = 0.923, intercept = 4.52; Figure 2B). Surprisingly, there was large disagreement between the results of older (Bismark and BWA meth) and newer methods (Biscuit and BiSulfite Bolt). Mean percent methylation per fish was much higher in Biscuit compared to BWA meth (slope = 0.377, intercept = 57.8, Figure 2C). While BiSulfite Bolt and Biscuit were more similar to each other than they were to BWA meth or Bismark, the lower slope and higher intercept indicate that average methylation was not as consistent between these two methods (slope = 0.793, intercept = 14.2, Figure 2D) as it was between BWA meth and Bismark.

**Figure 2:**
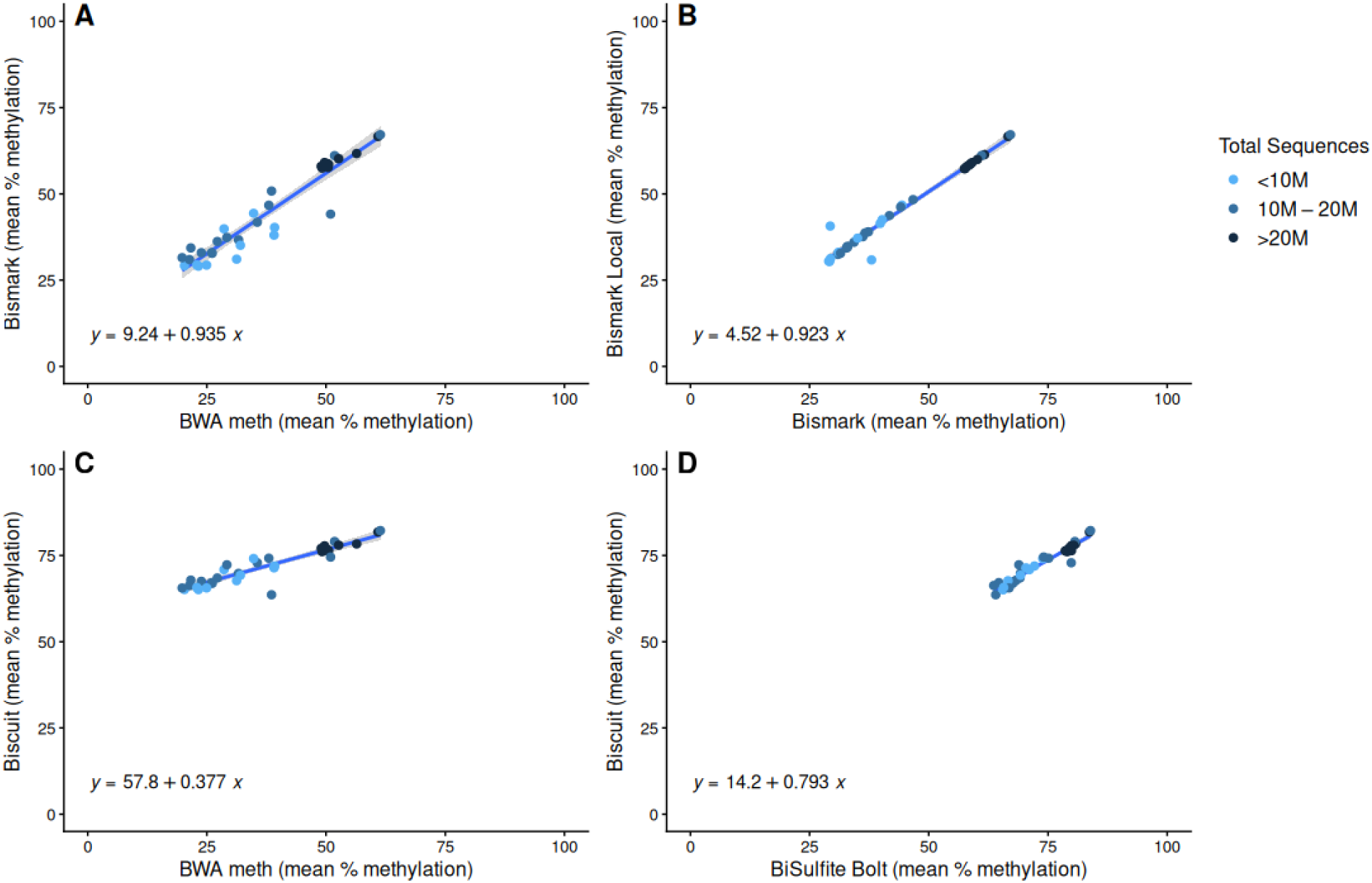
Mean percent methylation per base for individual stickleback sequenced with RRBS and analyzed using **(A)** BWA meth and Bismark, **(B)** Bismark global alignment and Bismark local alignment, **(C)** BWA meth and Biscuit, and **(D)** BiSulfite Bolt and Biscuit. A slope that is close to one and y-intercept near zero indicates agreement in mean percent methylation between read mapping methods.

We next leveraged the variation in sequencing effort among the stickleback sequencing batches to assess whether the total number of reads per sample influenced the reliability of methylation tabulation (Figure 3). With low sequence coverage there was wide variation between methods. An exponential decay model was a better predictor of the data than a linear model for all four methods comparisons. With the exception of the BWA meth-Biscuit residuals (Figure 3C), sequencing effort was a significant predictor of reduced residuals (BWA meth-Bismark: t = −3.061, p = 0.00444, Figure 3A; Bismark-Bismark Local: t = −2.084, p = 0.0455, Figure 3B; BiSulfite Bolt-Biscuit: t = −3.121, p = 0.00381, Figure 3D). Agreement in methylation calls increased with sequence coverage until approximately 20 million reads per sample, but increasing sequencing effort past this threshold provided little to no improvement (Figure 3).

**Figure 3:**
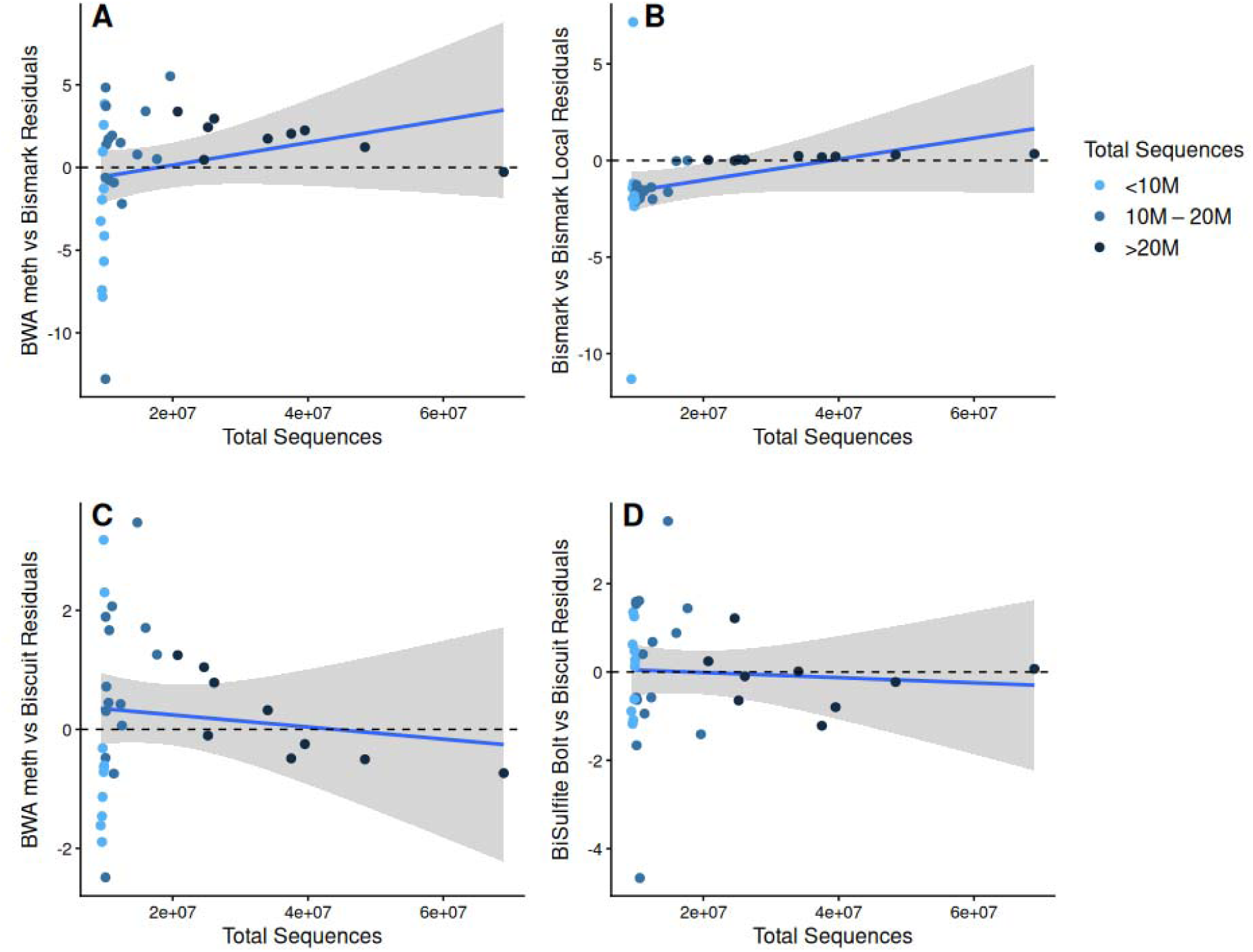
Total sequences per sample compared to the residuals (Figure 2) from models comparing **(A)** BWA meth and Bismark, **(B)** Bismark global versus Bismark local alignment, **(C)** BWA meth versus Biscuit, and **(D)** BiSulfite Bolt versus Biscuit.

We next examined distribution of percent methylation per base to further characterize how read mapping software influenced which CpG sites were retained for analysis (Figure 4, Supplemental Figures 4-6). Distributions obtained from older methods (Figure 4A-C) differed greatly from newer methods (Figure 4D-E), consistent with the previously noted differences in mean percent methylation obtained from each read mapping method (Figure 23). Notably, both Biscuit and BiSulfite Bolt recovered almost no unmethylated cytosines (percent methylation < 10%), while BWA meth, Bismark, and Bismark Local were dominated by unmethylated cytosines. Additionally, BWA meth, Bismark, and Bismark Local contained very few intermediately methylated sites (10-90% methylation).

**Figure 4:**
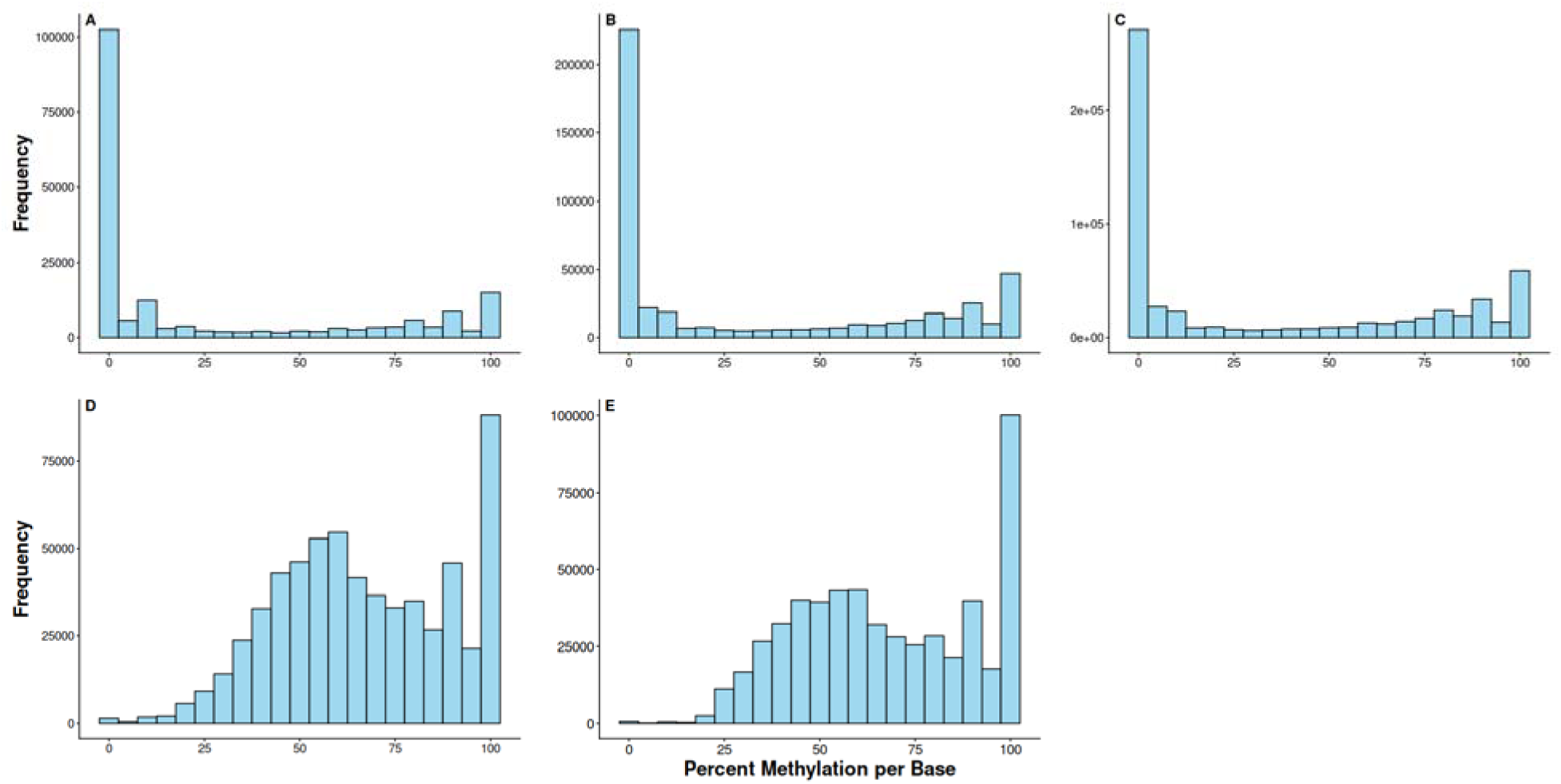
Frequency histograms of percent methylation per base for one RRBS sample from Wik lake (sample ID: CG_WK_002), analyzed with either **(A)** BWA meth, **(B)** Bismark, **(C)** Bismark Local, **(D)** Biscuit, or **(E)** BiSulfite Bolt.

### Differences in genomic breadth and depth of coverage between RRBS and WGBS

After running BWA meth and MethylDackel, the average median insert size of RRBS reads was 38.06 (± 2.44) compared to 242.25 (± 22.47) for WGBS. Despite the larger insert sizes and greater sequencing effort per sample for WGBS (RRBS batch 1: 10.5 ± 1.3M reads; RRBS batch 2: 31.5 ± 15.4M reads; WGBS: 58.5 ± 1.0M reads), RRBS still resulted in a higher per site depth (16.39 ± 3.46) than WGBS (12.73 ± 0.06).

We next compared the effects of applying different library preparation methods to technical replicates (i.e. RRBS vs WGBS; Supplemental Table 3). Applying a minimum depth filter of 10 reads per site on RRBS and WGBS libraries from the same individual resulted in ≤1.0% shared CpG sites. Lowering the minimum depth to 5 reads increased the proportion of shared sites to increase to 2.25% (± 0.42). Notably, at a minimum of 10x depth, >96% of CpG sites were exclusive to WGBS, highlighting the large difference in breadth of coverage between RRBS and WGBS (Supplemental Table 3).

When assessing CpG sites shared among biological replicates using RRBS, the percent of overlap depended on fish population. With a minimum of 10x depth, 13.1% and 27.2% of CpG sites were shared between individuals from either Watson or Wik Lakes, respectively. This population effect was much reduced with WGBS; 18.8% of CpG sites were shared between Watson Lake individuals and 19.7% between Wik individuals. Lowering the minimum depth to 5x increased the proportion of shared CpG sites using RRBS (28.4% for Watson, 40.1% for Wik) and WGBS (57.6% for Watson, 59.0% for Wik) (Supplemental Table 4).

Notably, the vast majority of the intermediately methylated CpG sites (i.e. the sites that are not always methylated or unmethylated), are not represented in the RRBS data (Figure 5). At a minimum depth of 10x, the proportion of CpG sites with methylation rates >10% and < 90% was nearly two-fold higher in WGBS (55.63 ± 2.49%, Figure 5A) than RRBS samples (28.56 ± 8.37%, Figure 5B). We recovered a higher proportion of intermediately methylated sites in batch 2 of RRBS sequencing (Supplemental Figure 7), but none of these samples were also analyzed via WGBS.

**Figure 5:**
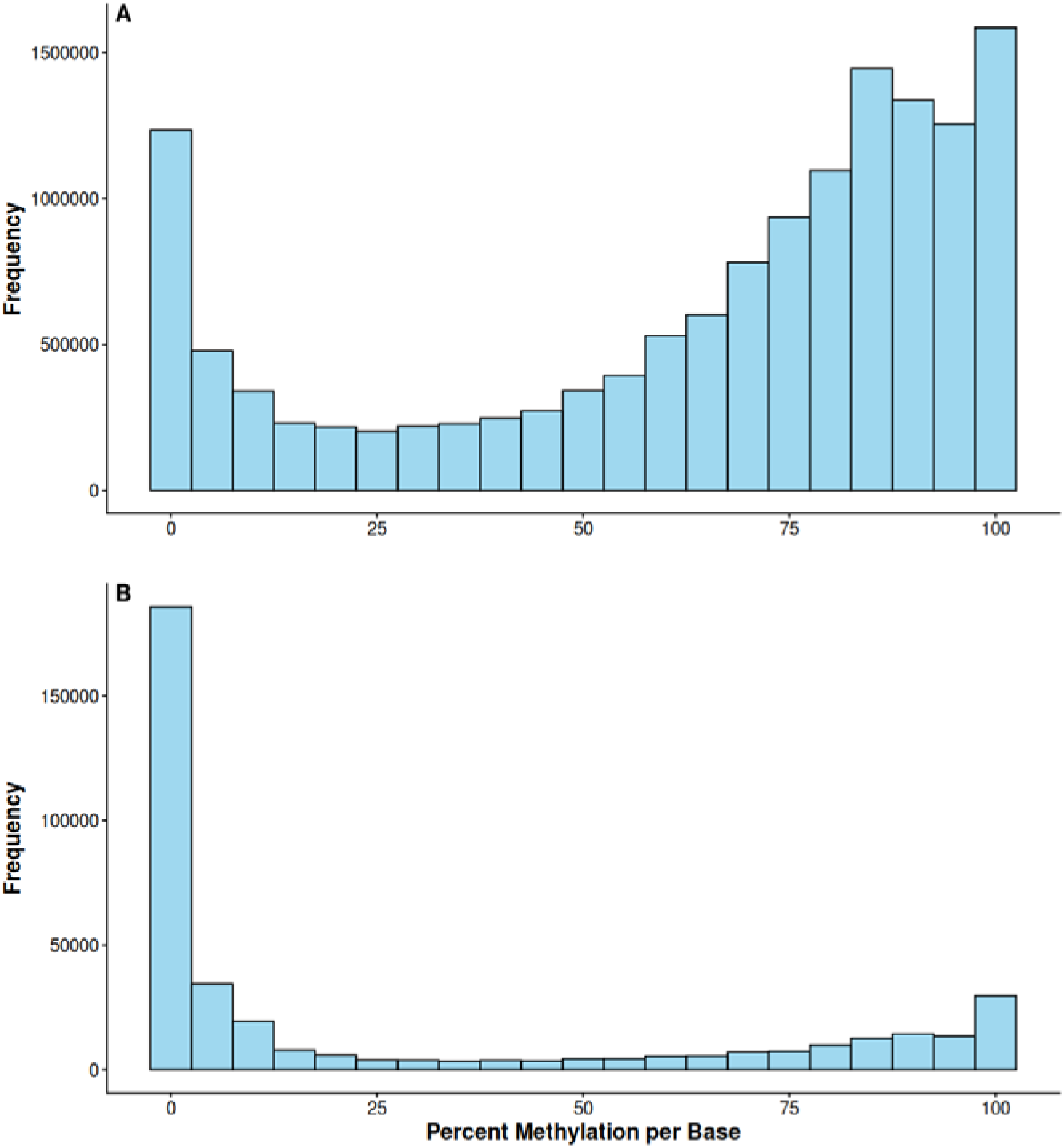
The percent methylation per base of sample CG_WK_002 sequenced with **(A)** WGBS and **(B)** RRBS after being read mapping with BWA meth.

Methodological differences in genomic breadth were also evident when assessing the distribution of CpG sites across functional categories. We conducted annotations using a minimum depth threshold of 5x to increase the number of shared CpG sites between library preparation methods, enhancing our ability to meaningfully compare the methods. CpG sites unique to WGBS recovered nearly all CpG islands, shores, and promoters, and 90% of exons and introns present in the threespine stickleback reference genome (Figure 6C & F). Additionally, the number of CpG sites available for functional annotations was 14.7x higher from sites unique to WGBS than sites unique to RRBS (Figure 6B & E). Because of the large difference in sequencing breadth, the intersection of both methods (i.e. CpG sites in common between WGBS and RRBS) reflects the same patterns found from sites recovered in RRBS only. However, CpG sites that were unique to RRBS still captured nearly 50% of CpG islands (Figure 6C) and 60% of promoters in the stickleback genome. Relatively few gene bodies were captured with RRBS (i.e., <15% of exons and introns; Figure 6F).

**Figure 6:**
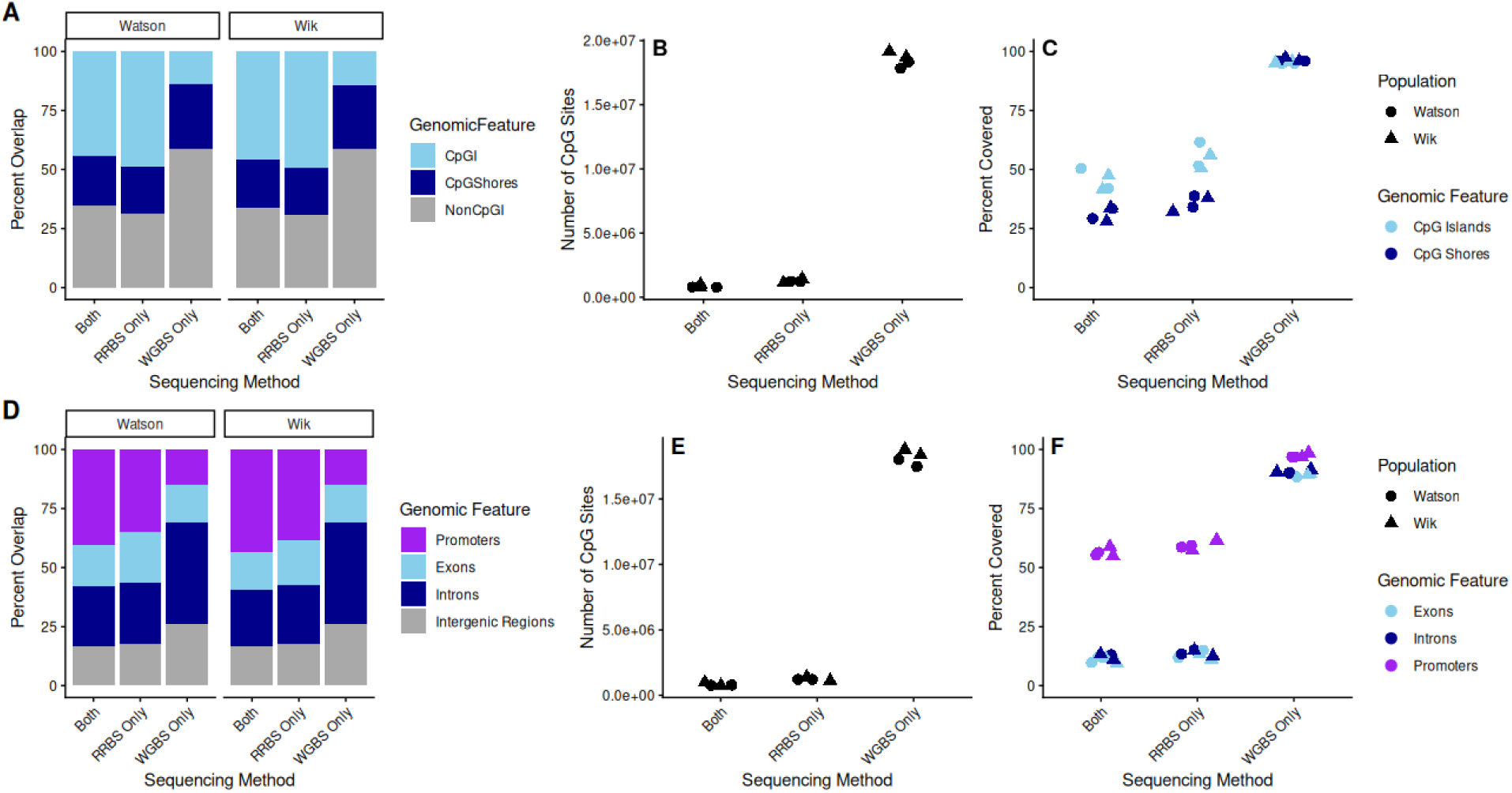
Comparisons of CpG sites that were shared or unique across sequencing methods, with all metrics averaged across replicates from each population.**(A)** Proportion of shared and unique CpG sites that overlap with CpG islands and shores. **(B)** Total number of observed sites. **(C)** Proportion of sampled CpG islands and shores in the threespine stickleback reference genome. **(D)** Proportion of sites that overlap with promoters, exons, introns, and intergenic regions. **(E)** Total number of CpG sites used for functional annotations. **(F)** Proportion of sampled promoters, exons, and introns in the threespine stickleback reference genome.

While WGBS clearly provides a larger breadth of coverage, RRBS enriches for functional genomic regions. When averaged across replicates from the same population (n=2), almost half of CpG sites unique to RRBS were found within CpG islands and 37% were within promoters. Meanwhile, a much smaller proportion of sites unique to WGBS overlapped with CpG islands (27%) and promoters (15%) (Figure 6A & D). Additionally, 26% of WGBS-only sites were in intergenic regions, compared to 17% of sites unique to RRBS. Overall, WGBS predominantly profiles CpG sites in intergenic regions or introns, while the majority of CpG sites sequenced by RRBS are in or near CpG islands, promoters, and exons.

### Concordance between RRBS and WGBS methylation calls

To assess agreement in methylation calling between RRBS and WGBS, we merged datasets from technical replicates by genomic position and calculated differences in percent methylation at each base obtained with WGBS and RRBS. We found broad agreement on a nucleotide-level between library preparation methods (RRBS percent methylation ∼ WGBS percent methylation: t = 2,467.4, p < 0.0001, R^2^ = 0.735). Almost 60% of CpG sites had ≤10% difference in percent methylation, which is within the margin of error according to the methylKit documentation (Figure 7A). As read depth influences methylation call accuracy, we also assessed the difference in methylation as a function of depth at each CpG site for both WGBS (Figure 7B) and RRBS (Figure 7C). We observed a significant but negligible relationship between depth and disagreement in percent methylation (R^2^ < 0.01).

**Figure 7:**
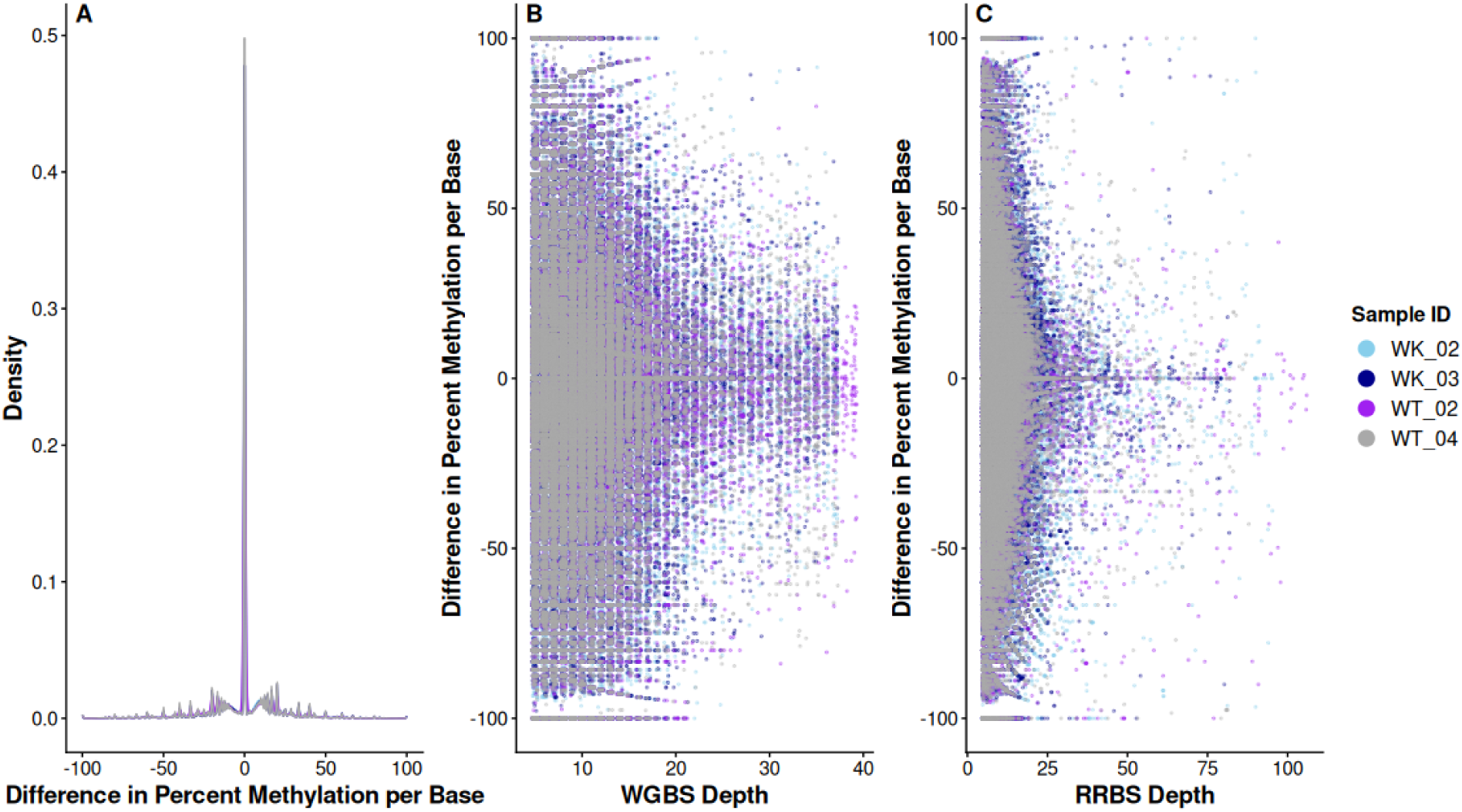
Comparisons of per site methylation estimated from different library preparation methods. **(A)** Difference in percent methylation per base obtained from RRBS and WGBS. Negative values indicate CpG sites with higher methylation in RRBS than WGBS. The relationship between the difference in percent methylation on a nucleotide level between library preparation methods and the depth of **(B)** WGBS and **(C)** RRBS data.

## Discussion

The increasing popularity of RRBS for methylation profiling has coincided with a rise in the use of Bismark as the default method for read mapping and methylation calling, particularly in ecologically focused experiments (Table 1). However, little effort has been dedicated to understanding the strengths and limitations of these methodological choices, particularly in genetically variable, wild organisms. By comparing RRBS and Bismark against less commonly used library preparation and analysis alternatives, we identify several options for researchers to maximize their sequencing resources.

### Optimizing bisulfite alignment software for ecological epigenetics

As predicted by previous studies (Farrell et al., 2021; Nunn et al., 2021; Zhou et al., 2024), BWA mem-based read mappers outperformed and Bismark in their ability to map the majority of sequenced reads to a reference genome (Figure 1). However, it is noteworthy that this previous work focused on human and plant data. We could find no previous studies assessing performance of bisulfite alignment wrappers across wild-caught/non-model systems. Despite differences in mapping efficiency, BWA meth and Bismark provided consistent estimates of the mean percent of CpG sites methylated per individual. However, there was a clear disagreement between more recent read mapping software and older software, in which Biscuit and BiSulfite Bolt estimated substantially higher methylation rates than BWA meth or Bismark (Figure 2). When we explored this trend further, we found that it was driven by a greater number of unmethylated cytosines captured by BWA meth and Bismark while Biscuit and BiSulfite Bolt recovered more intermediately methylated cytosines (Figure 4). Disagreement between read mappers could partially be alleviated through greater sequencing, but sequencing effort alone was insufficient to correct for large differences in software output (Figure 3).

For projects focusing on metrics related to per-site methylation, a key goal is to maximize the number of usable sequencing reads. Bismark resulted in the lowest mapping efficiency across all organisms and both BS-seq library types analyzed here, yet it remains the most commonly used read mapping method (Table 1). Continued reliance on Bismark may partly be attributed to it being one of the first bioinformatic tools designed to handle bisulfite converted data. Because Bismark does *in silico* conversions of all reads (sense and antisense, read and reference), its reliability is also agnostic to library directionality. Although Bismark’s ability to both map reads and call methylation levels makes it particularly accessible for researchers new to epigenetics, the tool’s default parameters may cause difficulties when working with paired end reads, as we observed when we initially had a 28% mapping efficiency for our stickleback RRBS data. Although rarely discussed in the literature, low mapping efficiency of paired end reads appears to be a relatively common issue with this tool (e.g., Nunn et al., 2021; Zhou et al., 2024). Indeed, the developers provide thorough documentation to help users resolve this mapping problem. The guidance highlights failure to trim adapters prior to read mapping, potential contaminants, and high stringency under default parameters, among other possible sources of error. Under default parameters, both paired end reads must map to one unique version of the *in silico* converted reference genome, otherwise the paired end sequence gets discarded. Following the advice in the documentation, and in-line with Nunn et al. (2021) and Zhou et al. (2024), we relaxed this threshold by mapping our reads as non-directional. This resulted in a mapping rate (54%) that is more in-line with previous DNAm studies in stickleback (Heckwolf et al., 2020; Hu et al., 2021). An additional complication with Bismark is the decision of whether to use the default global alignment method, in which sample reads must meet strict alignment requirements from beginning to end, or allow soft clipping of read ends that do not align as well to the reference genome as the rest of the insert (i.e. “local alignment”). The developers of Bismark argue that higher mapping efficiency obtained from local alignment is due to erroneously mapped reads. However, the other three BS-seq aligners tested here, including two that have been published within the last five years, rely on local alignment. Indeed, Zhou et al. (2024) found that Bismark not only had a lower mapping efficiency compared to other methods, but also resulted in the lowest rate of reads optimally mapped to the genome. Additionally, Farrell et al. (2021) used simulated data to determine Bismark had among the highest methylation call error rate, particularly for genetically variable data. While we cannot confirm the accuracy of local alignment because we used real BS-seq data, simply running Bismark as a local aligner resulted in significantly higher mapping efficiency across all organisms and sequencing methods.

Despite requiring additional tools for methylation calling, BWA meth resulted in significantly higher mapping efficiency than Bismark. While a previous study found that BWA meth had higher error rates than alternative tools (Farrell et al., 2021), this analysis was based on both directional and nondirectional libraries. It is plausible that the increased error rate was driven by strand bias rather than being a software limitation. Following BWA meth, we used MethylDackel to tabulate methylation and filter SNPs. Conventional wisdom is that RRBS libraries should be sequenced using single end reads, as paired end reads may lead to bias by repeatedly sequencing the same sites (Krueger et al., 2012). However, this bias can be reduced by normalizing the data based on median read depth, and if genetic polymorphisms are unknown, the overlapping nature of paired end reads can be exploited by software like MethylDackel to call SNPs. Additionally, using MethylDackel for SNP filtering reduces errors associated with SNPs falsely being considered unmethylated cytosines. In addition to facilitating SNP calling, paired end sequencing ensures high mappability in regions with many genomic repeats, which tend to be enriched in RRBS (Sun et al., 2018). Similar to most studies of ecological model organisms, we did not have genotype data for either the focal individuals or populations, and were thus unable to verify the accuracy of SNP filtering using MethylDackel. For a comprehensive assessment of SNP filtering tools for BS-seq data, see Lindner et al. (2022), although they do not include MethylDackel in their analysis.

We discovered that modern read mapping algorithms captured very few unmethylated cytosines, while older wrappers captured few intermediately methylated sites. There are a number of features inherent to RRBS that make it difficult to map to a reference genome. Enzymatic shearing during library preparation targets CpG-dense regions of the genome while bisulfite conversion increases the number of mismatches between the sample read and reference genome. These processes cause RRBS reads to be repetitive with small insert sizes, consequently making them particularly difficult to map accurately to a reference. It is possible that some algorithms repeatedly align reads to the same regions of the genome, while other algorithms spread the sample reads across the genome. Because we used real data, we are unable to determine which read mappers resulted in the most accurate methylation profiles. However, BWA meth and Bismark produced methylation profiles that are more analogous to the expected bimodal distribution of percent methylation, with most sites being either unmethylated (≤10% methylation) or fully methylated (≥90% methylated) (Akalin et al., 2012; Seiler Vellame et al., 2021; Sun et al., 2018). As far as we are aware, this type of bias between these methods has not been reported previously, highlighting how conventional methods optimized and tested only on inbred model organisms and/or simulated data may not be suitable for genetically variable, wild populations.

### RRBS and WGBS vary in breadth of coverage and methylation calls

In addition to assessing how bisulfite alignment wrappers performed with RRBS data, we examined how methylation profiles varied across technical and biological replicates using RRBS and WGBS. Few CpG sites were commonly sequenced between technical (Supplemental Table 3) or biological replicates (Supplemental Table 4). When comparing the methylation patterns recovered from technical replicates, the number of intermediately methylated sites (10-90%) was much lower in RRBS compared to WGBS (Figure 5). However, RRBS enriches for functional regions, while WGBS sequences across the entire genome (Figure 6). Although relatively few sites were shared across datasets, those that were shared often had similar estimates of methylation percentage (Figure 7).

One potential explanation for the lack of congruence between sites sequenced with RRBS and WGBS is that our sequencing parameters (i.e., 2×150bp reads) were much longer than the small inserts (∼38bp) produced via RRBS. This resulted in only a fraction of the total expected sequence data being useful for analyses. Importantly, increasing sequence effort led to greater agreement in downstream methylation calling, regardless of read mapping methods. However, this benefit plateaus at ∼20 million reads per sample (Figure 3). A previous study used simulations based on DNAm profiles from the cortex region of brains in an inbred mouse mouse strain to determine that the relationship between depth of coverage and methylation call accuracy increases rapidly until 10x depth of coverage, at which point the trend becomes asymptotic (Seiler Vellame et al., 2021). Comparing our stickleback results with those of Seiler Vellame et al. (2021) suggests that the optimal target coverage depth may vary among species, and also likely depends on what level of precision is important to investigators.

Interestingly, the level of concordance between biological replicates depended on both population and sequencing method. Individuals from Wik lake had over twice as many overlapping CpG sites compared to samples from Watson lake when sequenced using RRBS (Supplemental Table 4). Because this pattern was not present in WGBS, RRBS library preparation is the likely culprit. Despite having a similar number of reads, Watson fish had fewer CpG sites and a slightly higher read depth (14x vs 13x) than individuals from Wik. Taken together, this suggests that fewer MspI cut sites (or a decrease in enzymatic cleavages) in Watson fish led to less CpG sites following library preparation, resulting in the sites that were recovered to be sequenced at a slightly higher depth. To test this prediction, we would need population-specific genomes for *in silico* digestion and a larger sample size than n = 2/population. These types of data limitations are common in studies of natural populations. Overall, these results highlight the importance of assessing common methods across a diversity of taxa and populations, as genetic variation can affect what data is available for downstream analysis.

When assessing methylation profiles of WGBS-RRBS technical replicates, the proportion of intermediately methylated sites sequenced with WGBS was almost twice as high as intermediate methylation captured with RRBS (56% versus 29%, respectively). Methylation can degrade if tissue samples are collected multiple hours after mortality (Rhein et al., 2015) or frozen for a long period of time (Lee et al., 2023), but the liver tissue we sequenced was stored in RNALater immediately upon euthanasia and kept at −70 for less than one year. Therefore, degradation of intermediately methylated sites is unlikely the main driver of this observation. Rather, there are technical and evolutionary explanations that likely interacted to create this pattern. Batch 2 RRBS samples recovered a much higher proportion of intermediately methylated sites than batch 1 (Supplemental Figure 7), likely driven by more sequencing. However, this higher rate (37.83 ± 5.78%) is still substantially lower than the average of WGBS, suggesting that there may be fundamental, biological differences in the methylation patterns surveyed by each method. Indeed, while WGBS captured almost all of the CpG islands, promoters, introns, and exons in the stickleback reference genome, over half of WGBS CpG sites were found in introns or intergenic regions. Meanwhile, over half of the CpG sites unique to RRBS were in promoters and exons. Given that molecular ecologists are likely interested in identifying functionally important epigenetic effects, RRBS may provide a better option for detecting and measuring these changes.

We found consensus when comparing methylation rates on a single base pair level between technical replicates sequenced with RRBS and WGBS, confirming the validity of both library preparation methods for assessing DNAm. This concordance would likely have been higher if we used a more stringent SNP filter. Our SNP filtering cut off was rather lenient, meaning that we likely called some sites as methylated cytosines when in reality they were due to genetic variation. When RRBS and WGBS disagreed on methylation calls, it was equally common for RRBS to call a site as being more or less methylated than WGBS, suggesting that there is not a methodological bias. However, bisulfite conversion damages DNA at unmethylated cytosines during library preparation, which inherently inflates genome-wide methylation levels (Olova et al., 2018). The magnitude of this bias is only recently becoming clear thanks to newer library preparation approaches that forego harsh chemical reactions and instead rely on enzymes to convert unmethylated cytosines (e.g., EM-Seq: Vaisvila et al., 2021). If precise estimates of genomewide methylation are of key concern, then consideration should be given to using these newer, non-bisulfite methods of library preparation. Importantly, EM-seq converts unmethylated cytosines to thymine, making the sequencing data compatible with existing bioinformatic tools for bisulfite converted data. Despite these advances, RRBS remains the most cost effective DNAm profiling method and is likely sufficient if the primary goal of a project is to measure or detect relative changes in methylation (i.e., those likely to have functional effects or are environmentally-induced). However, more studies should be done to assess agreement between technical replicates sequenced with WGBS, RRBS, and EM-seq to validate these methods across a variety of organisms.

## Conclusions

We take a critical approach to analyzing RRBS and WGBS data using several conventional methods. We highlight the strengths and weaknesses of both molecular and bioinformatic methods, all of which should be considered during experimental design. If SNP filtering in the absence of genotype data is a priority, researchers may consider choosing paired end sequencing, contrary to traditional advice regarding RRBS library construction. Given that RRBS can result in exceedingly small insert sizes and there may be random variation in the amount of sequencing effort for each sample, we suggest using relatively short read lengths and deeply sequencing a small number of initial samples to determine the sequencing threshold required to obtain accurate per site methylation metrics. In the case of threespine stickleback, 20 million reads/sample appears to be the optimal trade off between cost and accuracy. Additionally, the most commonly used tool for analyzing BS-seq data (Bismark) performs significantly worse than alternative methods in terms of percentage of reads mapped. However, more work needs to be done to reconcile the difference between methylation profiles obtained using older and newer read mapping software. Finally, we highlight how different library preparation methods recover distinct methylation profiles, and why researchers may consider alternative library preparation methods to BS-seq.

## Acknowledgements

We thank Sean Schoville for computing resources, and Kristofer Sasser and Cole Wolf for collecting fish from Vancouver Island. We recognize that the fish populations analyzed here originate from lakes in the unceded territory of the Dena’ina and Coast Salish people and thank them for their continued land stewardship. This work was funded by the Department of Integrative Biology at UW-Madison and National Institute of General Medical Sciences 5R35GM142891-02.

## Data Accessibility and Benefit-Sharing

*Data Accessibility Statement:* All scripts are available at https://github.com/EmilyKerns/VariablePerformanceofBS-SeqandReadMappingSoftware. Stickleback sequence data is available on Dryad at https://doi.org/10.5061/dryad.dncjsxm9h and NCBI SRA (BioProject Accession Number PRJNA1439918).

## Author Contributions

E.V.K. and J.N.W. conceived and designed the study and co-wrote the manuscript; E.V.K. wrote the scripts, conducted analyses, and produced figures; J.N.W. supervised and funded the work.

## Ethics Approval Statement

All animals were handled and euthanized in accordance with UW-Madison IACUC protocol L006460-A01. Field sampling was done under the Alaska Department of Fish & Game permit number SF2022-043 and BC Ministry of Forests permit number NA23-787881.

