## Supplemental Information for "Variable performance of widely used bisulfite sequencing methods and read mapping software for DNA methylation"

### **Supplemental Methods:**

#### *Literature review of bisulfite library preparation and analysis methods*

To assess how frequently Bismark, BWA meth, BiSulfite Bolt, Biscuit, and MethylDackel are used to analyze methylation data, we used “software name” AND “DNA methylation” to query Web of Science and Google Scholar databases on September 8, 2025. We consolidated the number of search results from each database in Table 1.

*Sample and Data Collection*

Fish were euthanized with an overdose of buffered MS-222 according to IACUC protocol L006460-A01. Field samples were dissected lakeside to avoid introducing transportation stress that may influence DNAm profiles (Alaska Department of Fish & Game permit number SF2022-043, BC Ministry of Forests permit number NA23-787881). We dissected and stored liver tissue in RNALater (Sigma-Aldrich). We chose liver as our focal tissue because previous work suggests liver-specific methylation is relatively conserved across vertebrates (Klughammer et al., 2023). Liver also has relatively few cell types (Gumucio & Miller, 1981), which should limit between sample variation due to cell type specific methylation. Two lab fish from each of the Roberts, Gosling, and Sayward populations were exposed to the tapeworm *Schistocephalus solidus* as part of a separate study, and one of the Sayward individuals was successfully infected. None of these individuals displayed obvious differences in DNAm patterns, and therefore were retained for further analysis (Supplemental Figure 1).

#### *Threespine Stickleback Sequencing*

We extracted DNA from each stickleback liver sample using carboxyl coated magnetic beads (BOMB.bio method: Oberacker et al., 2019). Bisulfite conversion, library preparation, and sequencing were performed by Admera Health Biopharm Services. RRBS samples were bisulfite treated and then converted to Illumina sequencing libraries and using Zymo-Seq RRBS Library Kit. WGBS were similarly bisulfite converted using EZ DNA Methylation-Gold Kit (Zymo Research, California, USA) and library preparation was done with xGen™ Methylation-Sequencing DNA Library Preparation Kit (Integrated DNA Technologies, California, USA). Sample quality was assessed using Quibit, agarose gel, and qPCR prior to sequencing. Libraries were sequenced on Illumina Novaseq 6000 using paired end 150bp read chemistry. DNA extractions and sequencing were performed in two batches. Each barcoded RRBS sample was mixed into one of two sequencing pools (Supplemental Table 1). All WGBS samples were sequenced in one batch.

#### *Statistical analyses of BS-seq Alignment Wrappers*

For RRBS stickleback samples, which received relatively low or high sequence coverage depending on batch, we used Wilcox rank sum exact tests to test whether sequencing batch affected mapping efficiency for each of the read mapping tools (Supplemental Figure 3). To determine if the resulting methylation profiles of individuals were in agreement despite differences in read mapping approach, we used methylKit v. 1.33.3 (Akalin et al., 2012) to calculate the mean percent methylation for each sample. We then created a linear model in ggpmisc v. 0.6.1 (Aphalo, 2024) to compare the mean percent methylation obtained using each pipeline. To assess whether variation in the amount of sequencing effort for each sample may explain inconsistencies in mean percent methylation, we visualized the relationship between the residuals of each model and the total number of sequences for each sample. The association between the absolute value of the residuals and total sequences per sample was tested by comparing the fit of a linear and exponential decay model using AIC scores.

#### *Genomic Annotations of RRBS-WGBS Technical Replicates*

We next assessed differences between RRBS and WGBS in the threespine stickleback. To determine if average read depth per sample differed between library types we used a Wilcox rank sum test. To determine where RRBS and WGBS vary in their coverage of the genome, we conducted genomic annotations on CpG sites that were sequenced with both methods, RRBS only, and WGBS only with a minimum depth of 5x. We converted the methylation dataframes into GRanges objects (GenomicRanges v. 1.54.1) (Lawrence et al., 2013) and quantified the overlap between methylation sites and CpG islands and shores (±2 kbp CpG islands) with genomation v. 1.34.0 (Akalin et al., 2015). We next imported the annotated stickleback v.5 reference genome (Nath et al., 2021) using methylKit v. 1.27.1 to assess differences in CpG sites captured in both BS-seq methods, RRBS only, and WGBS only.

### **Supplemental Tables:**

**Supplemental Table 1:** Stickleback population, rearing environment, and sample sizes with associated batch of sequencing for RRBS and WGBS. Abbreviations for populations and rearing environments are consistent with figures in the results.

| **Population** | **Rearing Environment** | **Sample Size *(Sequencing Batch)*** |
| --- | --- | --- |
| Watson (WT, 60.54, -150.47) | Field (FD) | n=4 *(1)* |
|  | Common Garden (CG) | n=4 *(1)* |
|  |  | WGBS (n=2) *(1)* |
| Wik (WK, 60.72, -151.25) | Field | n=4 *(1)* |
|  | Common Garden | n=4 *(1)* |
|  |  | WGBS (n=2) *(1)* |
| Roberts (ROB, 50.22, -125.54) | Field | n=3 *(2)* |
|  | Common Garden | n= 2 *(1)*, n = 2 *(2)* |
| Gosling (GOS, 50.05, -125.50) | Field | n=3 *(2)* |
|  | Common Garden | n=2 *(1)*, n = 2 *(2)* |
| Sayward (SAY, 50.39, -125.95) | Common Garden | n=2 *(1)*, n = 2 *(2)* |

**Supplemental Table 2**: NCBI project number for each dataset used for analysis and the reference genome we used to analyze each dataset.

| **Organism** | **NCBI BioProject Accession Number** | **Reference Genome** | **GC% of the Reference Genome** | **Sample size** |
| --- | --- | --- | --- | --- |
| Threespine stickleback (*Gasterosteus aculeatus*) | PRJNA1439918 | GCF_016920845.1  (Nath et al., 2021) | 44.5% | RRBS n = 34,  WGBS n = 4 |
| Cichlid (*Astatotilapia calliptera*) | PRJNA728457 | GCF_000238955.4  (Brawand et al., 2014; Conte et al., 2015) | 41% | RRBS n = 33,  WGBS n = 6 |
| Great tit (*Parus major*) | PRJNA208335 | GCF_001522545.3  (Laine et al., 2016; Laine et al., 2019) | 41.5% | RRBS n = 61 |
| Purple sea urchin (*Strongylocentrotus purpuratus*) | PRJNA548926 | GCF_000002235.5  (Sea Urchin Genome Sequencing Consortium et al., 2006) | 37.5% | RRBS n = 12 |
| Coral (*Acropora nana*) | PRJNA1074434 | GCF_013753865.1  (Fuller et al., 2020) | 39% | WGBS n = 30 |
| Stick insect (*Timema cristinae*) | PRJNA1010130 | GCA_050494535.1  (Gompert et al., 2025) | 35.5% | WGBS n = 7 |

**Supplemental Table 3:** CpG sites that were in common or unique to WGBS and RRBS for technical replicates with a minimum of 10x depth and 5x depth threshold.

| Sample ID | Minimum Depth Threshold | Unique to WGBS *(Number of CpG sites)* | Unique to RRBS *(Number of CpG sites)* | Overlapping Sites *(Number of CpG sites)* |
| --- | --- | --- | --- | --- |
| CG_WK_002 | 10x | 96.6% *(5,417,478)* | 2.4% *(134,797)* | 1.0% *(55,001)* |
|  | 5x | 96.0% *(18,501,995)* | 1.1% *(219,188)* | 2.8% *(546,037)* |
| CG_WK_003 | 10x | 97.1% *(5,160,689)* | 2.2% *(116,016)* | 0.7% *(36,879)* |
|  | 5x | 96.9% *(18,194,034)* | 1.0% *(188,716)* | 2.1% *(402,195)* |
| CG_WT_002 | 10x | 98.8% *(4,966,649)* | 0.9% *(46,124)* | 0.3% *(14,544)* |
|  | 5x | 97.1% *(17,915,424)* | 1.1% *(194,016)* | 1.8% *(337,685)* |
| CG_WT_004 | 10x | 97.1% *(4,410,125)* | 2.3% *(102,397)* | 0.7% *(31,069)* |
|  | 5x | 96.5% *(17,312,003)* | 1.2% *(209,426)* | 2.3% *(418,643)* |

**Supplemental Table 4:** CpG sites that were in common between biological replicates from the same population using RRBS and WGBS with a minimum of 10x and 5x depth threshold.

| Sample Comparison | Sequencing Method | Minimum Depth Threshold | Unique to First Sample *(Number of CpG sites)* | Unique to Second Sample *(Number of CpG sites)* | Overlapping Sites *(Number of CpG sites)* |
| --- | --- | --- | --- | --- | --- |
| CG_WT_002 & CG_WT_004 | RRBS | 10x | 22.2% *(38,133)* | 64.6% *(110,931)* | 13.1% *(22,535)* |
|  |  | 5x | 30.4% *(274,939)* | 41.1% *(371,307)* | 28.4% *(256,762)* |
|  | WGBS | 10x | 44.0% *(3,493,419)* | 37.2% *(2,953,420)* | 18.8% *(1,487,774)* |
|  |  | 5x | 22.3% *(5,099,302)* | 20.0% *(4,576,839)* | 57.6% *(13,153,807)* |
| CG_WK_002 & CG_WK_003 | RRBS | 10x | 43.2% *(116,469)* | 29.5% *(79,556)* | 27.2% *(73,329)* |
|  |  | 5x | 39.0% *(377,396)* | 21.0% *(203,082)* | 40.1% *(387,829)* |
|  | WGBS | 10x | 41.7% *(3,715,901)* | 38.6% *(3,440,990)* | 19.7% *(1,756,578)* |
|  |  | 5x | 21.5% *(5,085,066)* | 19.6% *(4,633,263)* | 59.0% *(13,962,966)* |

### **Supplemental Figures:**

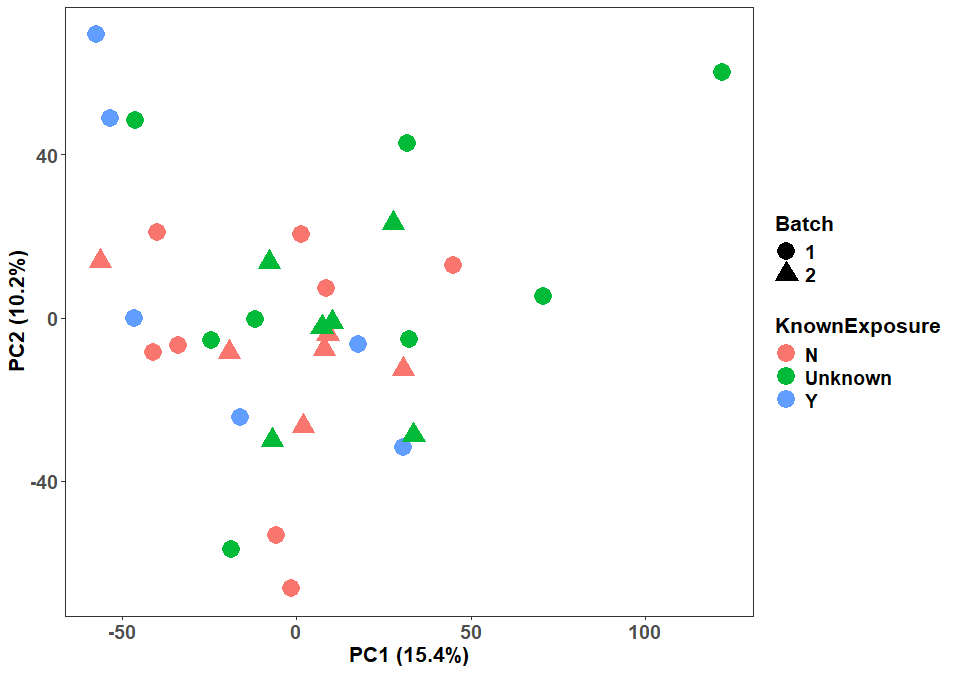

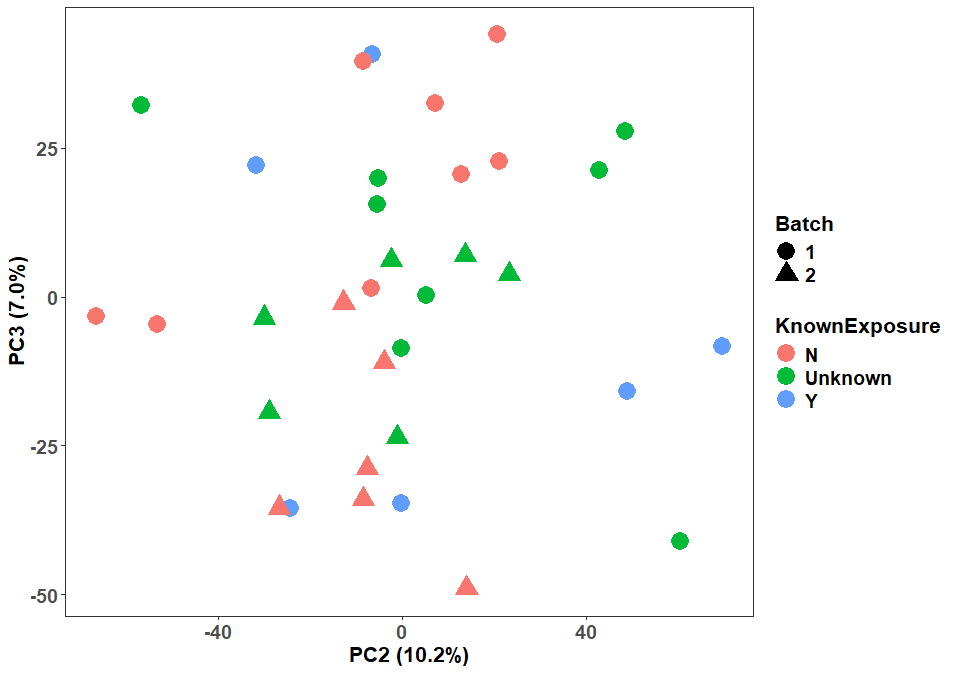

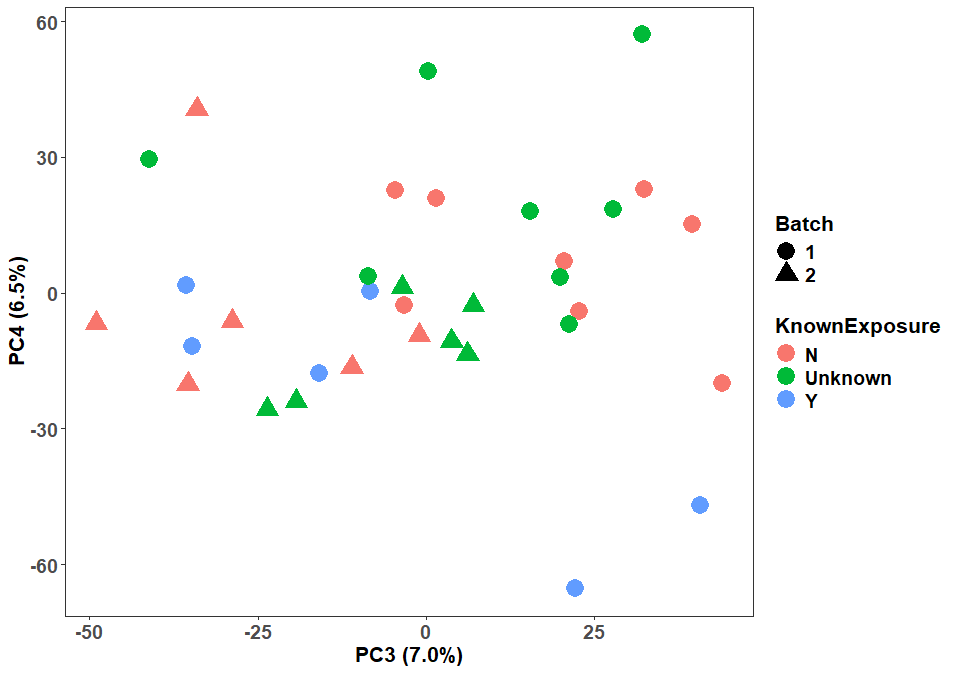

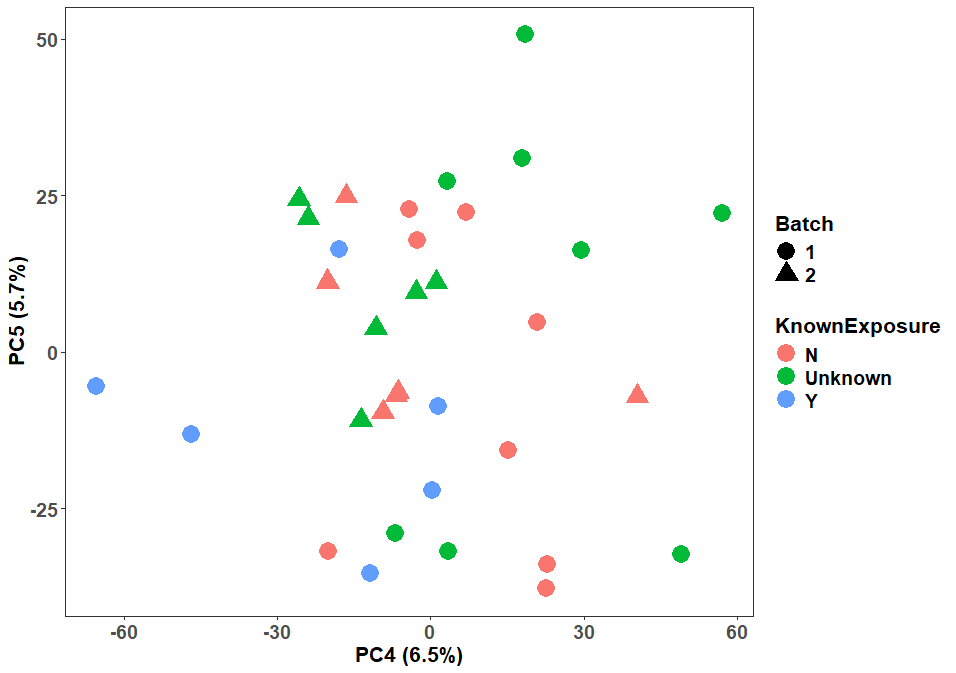

**Supplemental Figure 1:** PCA did not reveal any batch effects of known *S. solidus* exposure along PCs 1-5. All wild fish were considered “unknown.”

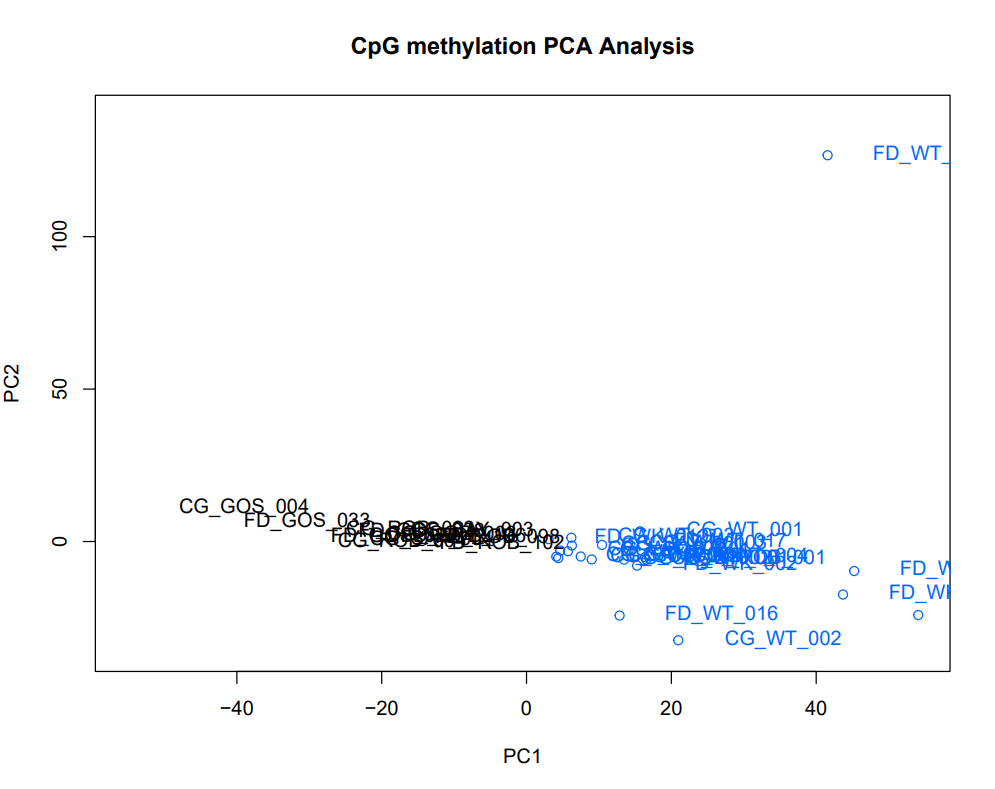

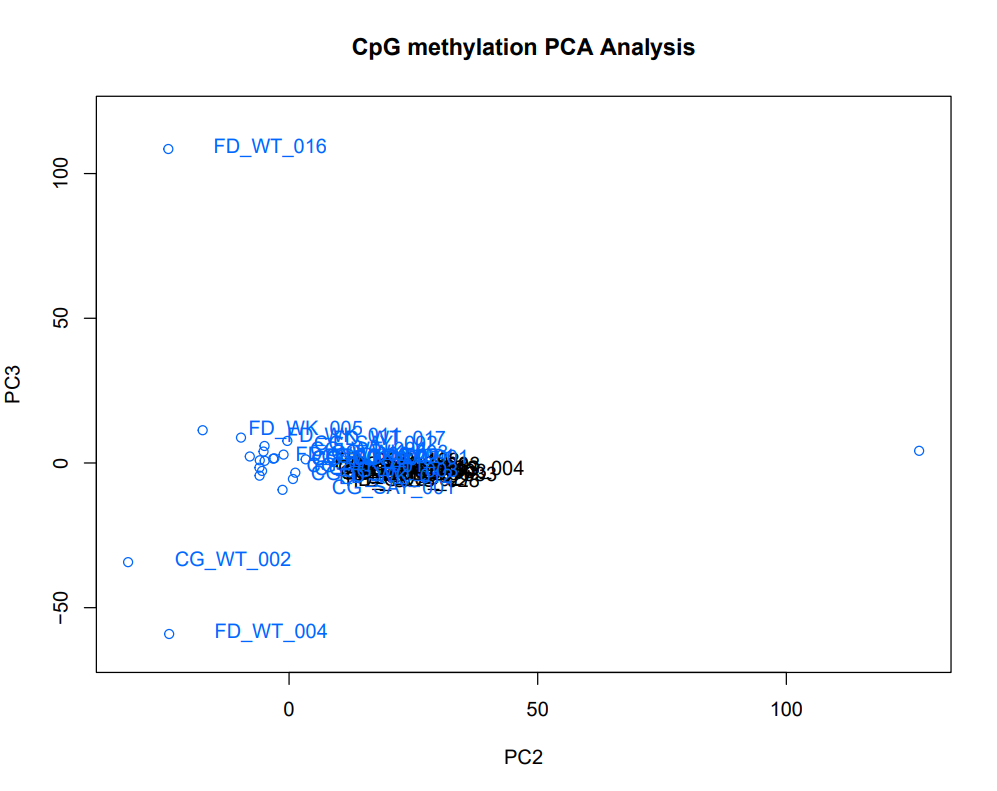

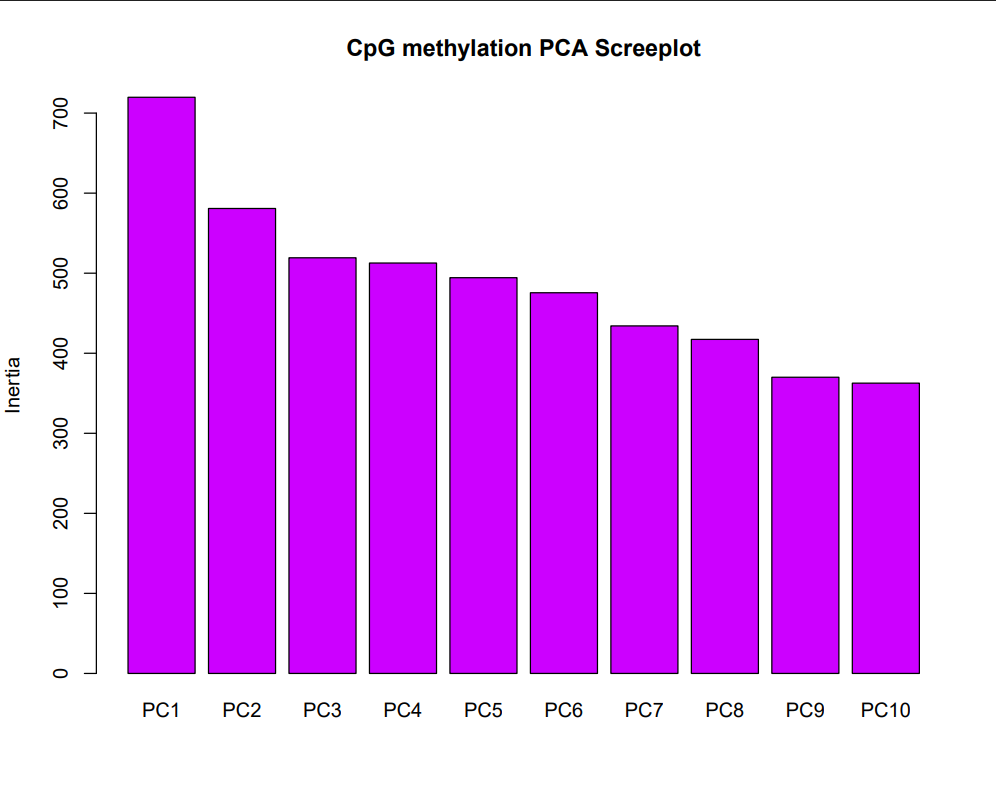

**Supplemental Figure 2:** Sequencing batch effects explained most of the variance in the data along PC1, but dissipated along PC2. Sample IDs in blue represent sequencing batch 1 while sample IDs in black represent sequencing batch 2.

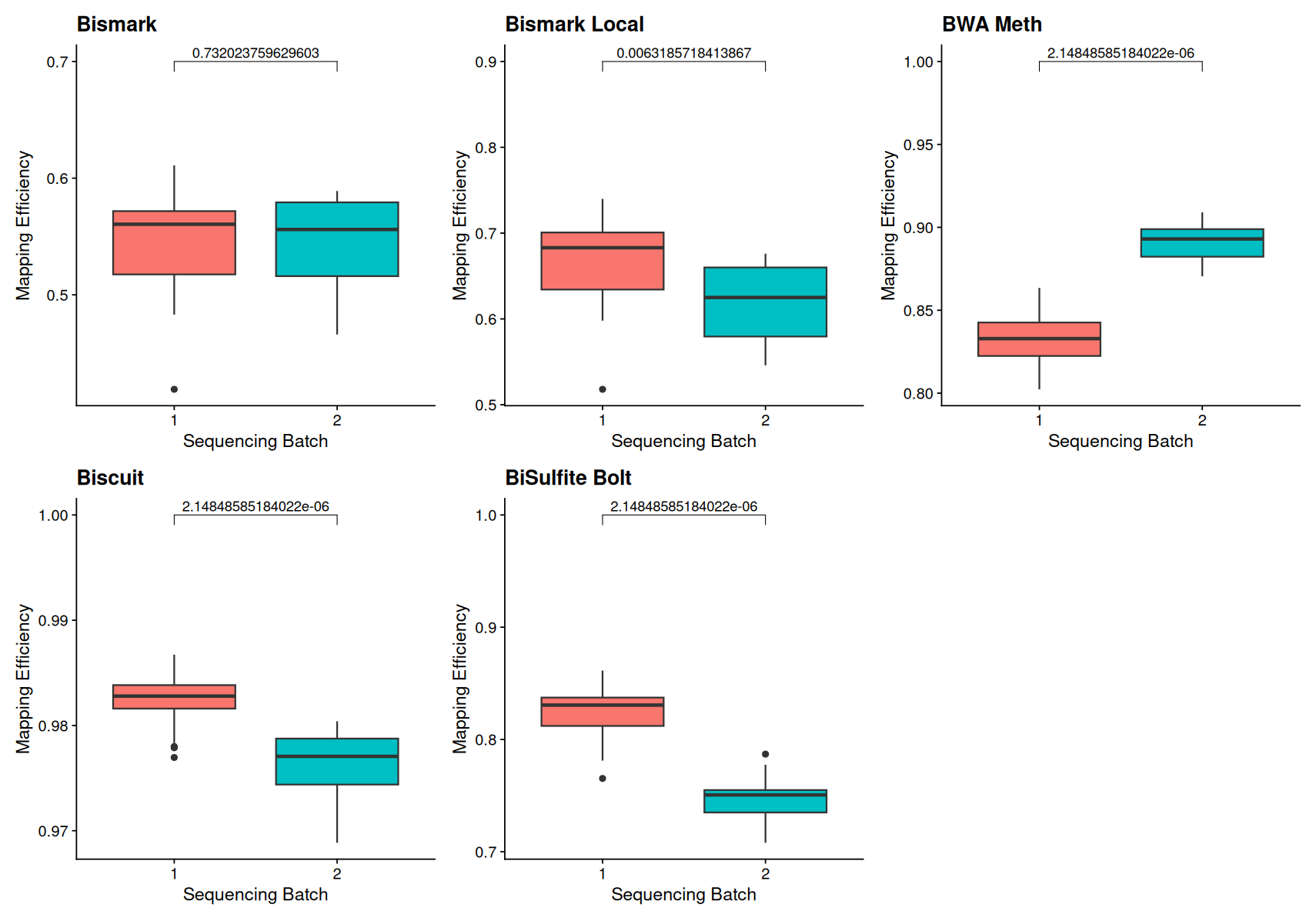

**Supplemental Figure 3:** All read mapping methods resulted in significant batch effects on mapping efficiency, except for Bismark (end-to-end alignment). The p value is above the lines that denote significance.

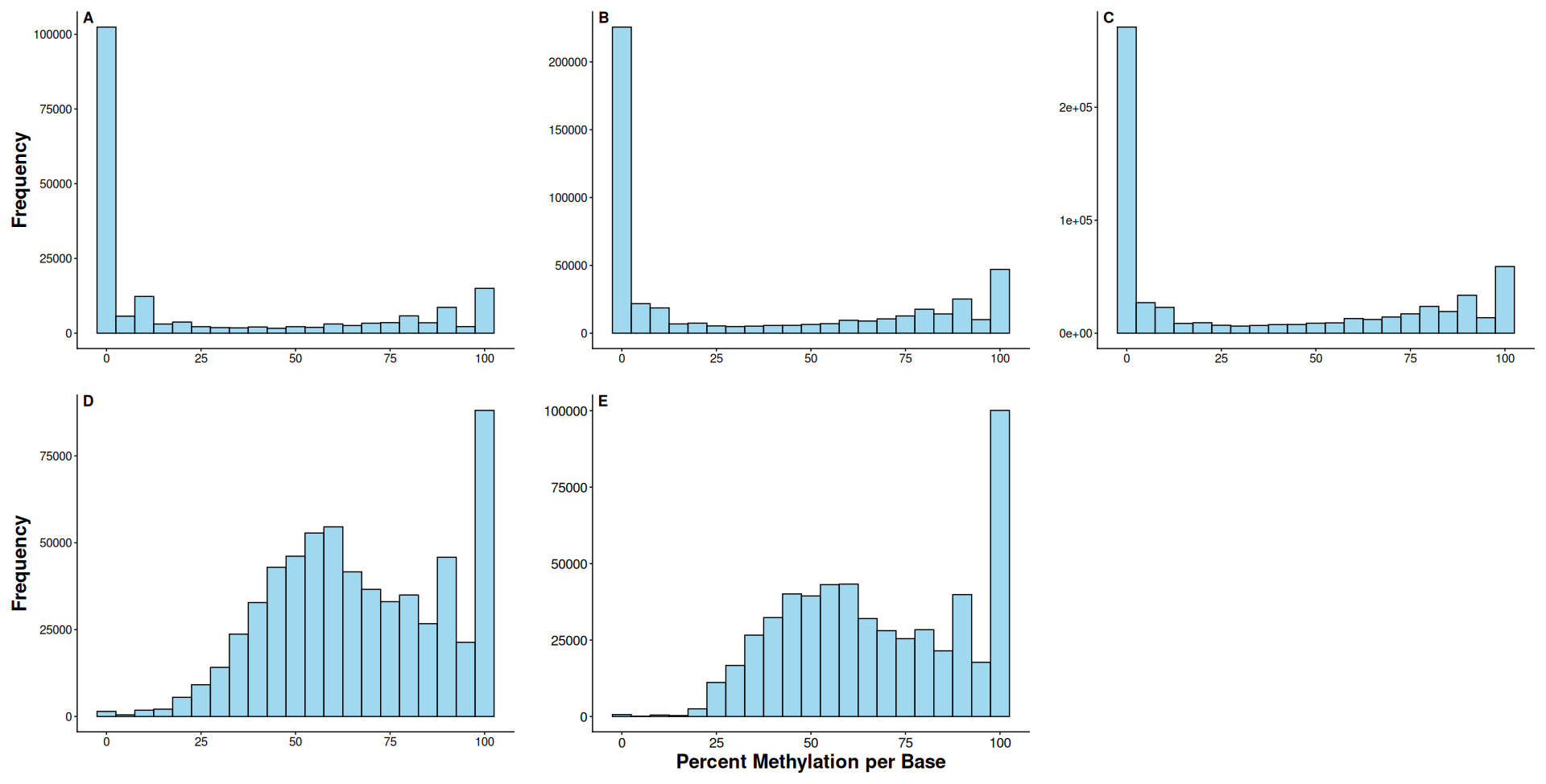

**Supplemental Figure 4:** Frequency histograms of percent methylation per base for a sample from Wik lake (sample ID: CG_WK_003). Liver tissue from this sample was sequenced using RRBS before read mapping with **(A)** BWA meth, **(B)** Bismark, **(C)** Bismark Local, **(D)** Biscuit, and **(E)** BiSulfite Bolt.

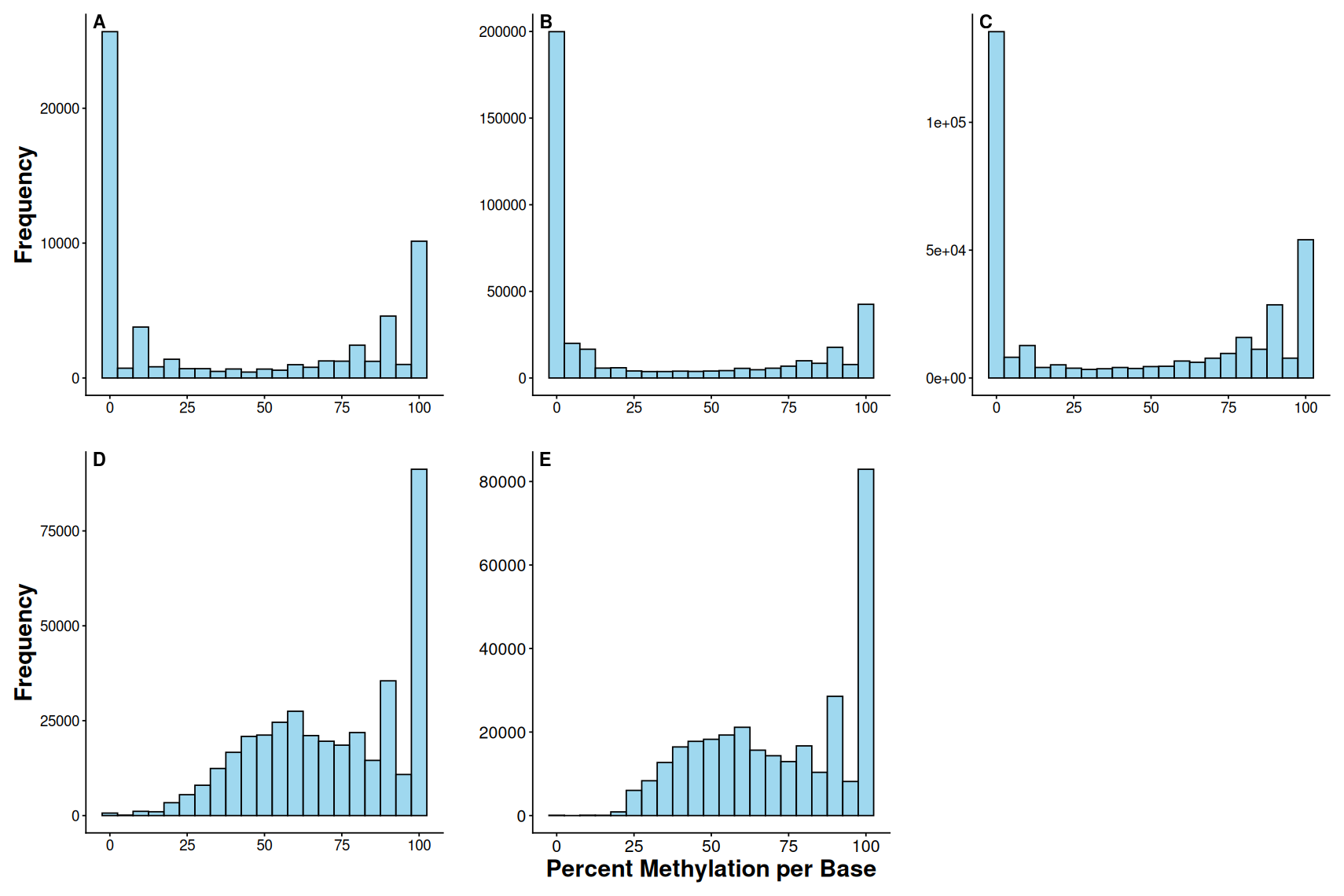

**Supplemental Figure 5:** Frequency histograms of percent methylation per base for a sample from Wik lake (sample ID: CG_WT_002). Liver tissue from this sample was sequenced using RRBS before read mapping with **(A)** BWA meth, **(B)** Bismark, **(C)** Bismark Local, **(D)** Biscuit, and **(E)** BiSulfite Bolt.

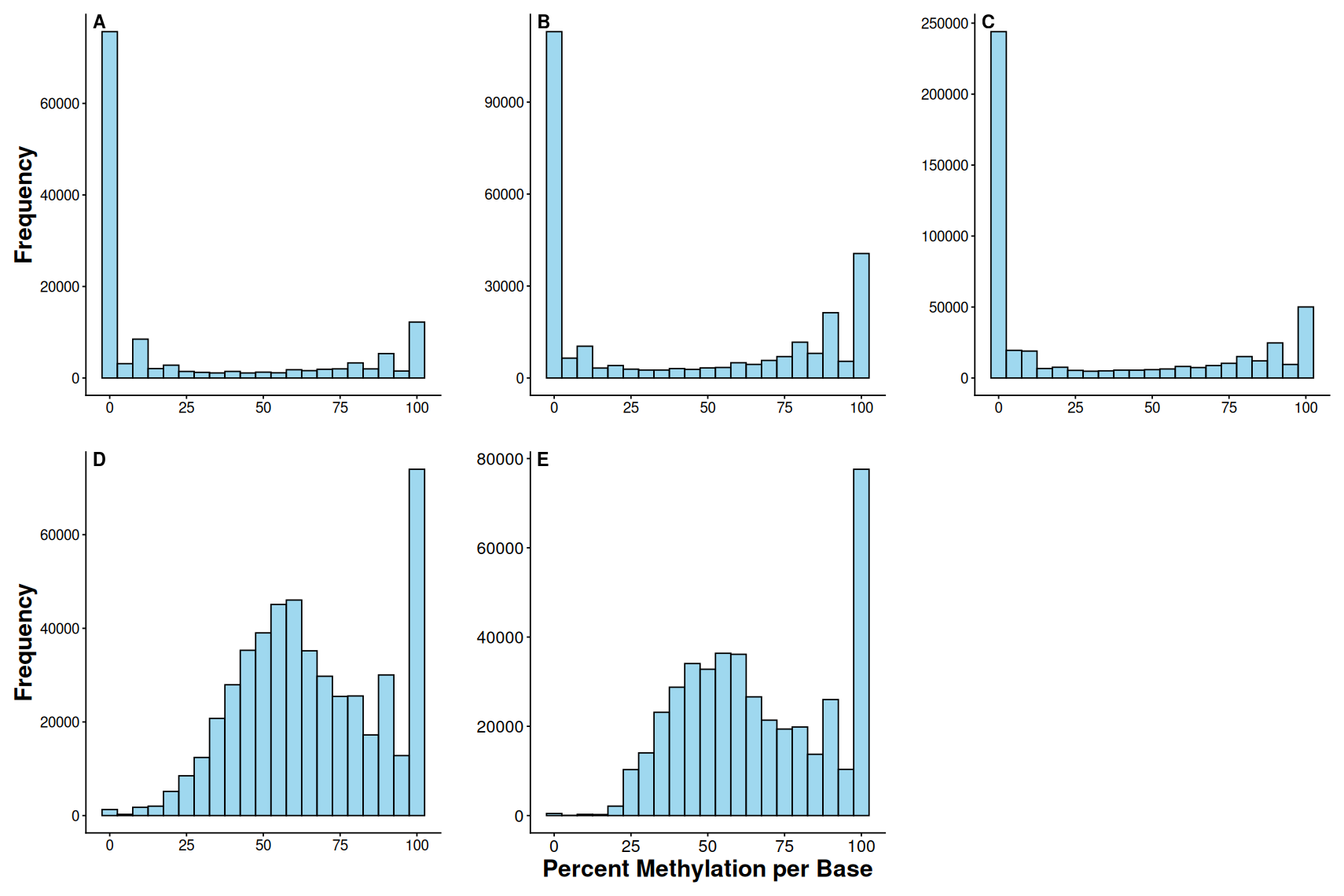

**Supplemental Figure 6:** Frequency histograms of percent methylation per base for a sample from Wik lake (sample ID: CG_WT_004). Liver tissue from this sample was sequenced using RRBS before read mapping with **(A)** BWA meth, **(B)** Bismark, **(C)** Bismark Local, **(D)** Biscuit, and **(E)** BiSulfite Bolt.

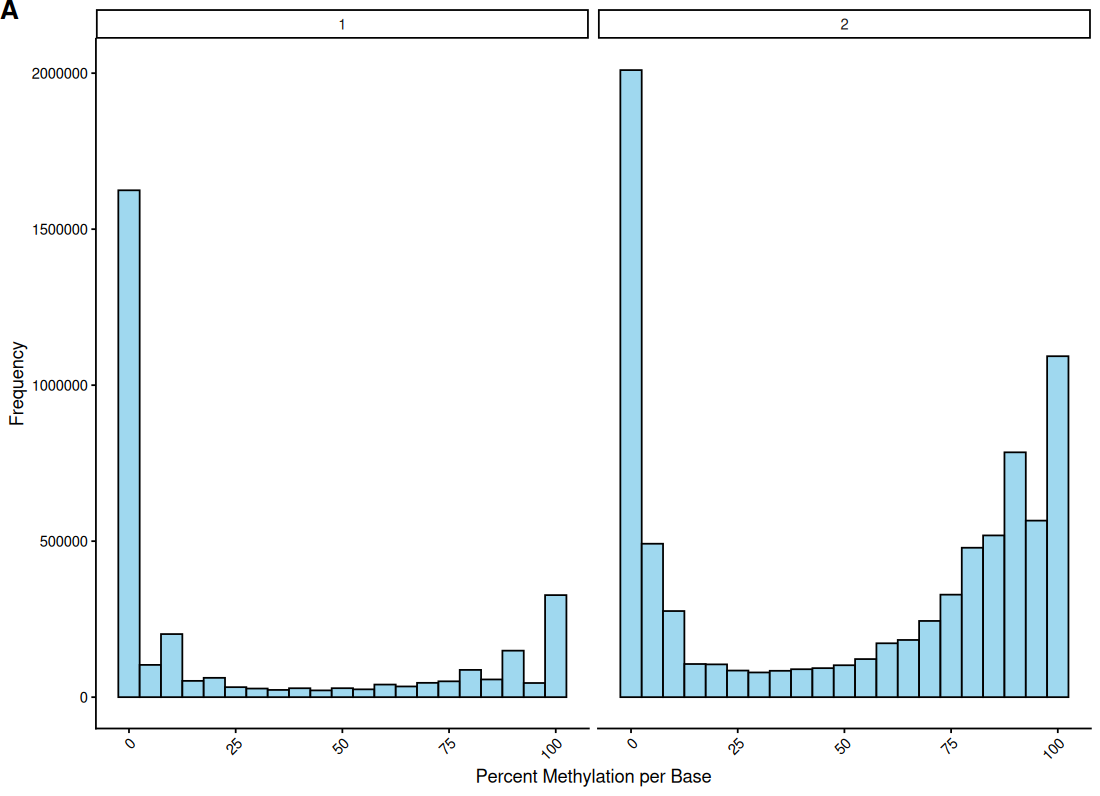

**Supplemental Figure 7:** Frequency histograms of percent methylation per site for RRBS samples based on sequencing batch. Batch 2 recovered more intermediately methylated cytosines (10-90% methylation) than batch 1, but only batch 1 had samples used as technical replicates for WGBS.
